# Volumetric building blocks: A simple and modular approach to create perfusable models via assembly of volumetrically printed subunits

**DOI:** 10.64898/2026.11.10.765611

**Authors:** Michael Bartolf-Kopp, Isabel Geroneit, Csaba Gergely, Franz Moser, Lisa Galaba, Nathaly Chicaiza Cabezas, Vivien Priebe, Marco Beaumont, Tomasz Jüngst

## Abstract

Tomographic volumetric additive manufacturing (tVAM) enables rapid 3D printing of constructs with intricate, freestanding geometries. Besides demonstrating very promising applications in tissue engineering and biofabrication, a limitation of the technology has been the relatively high material consumption and limited maximal model size due to optical and technical constraints. Here we demonstrate the fabrication and use of highly modular hollow structures created with tVAM to produce large scale tubular systems that allow the formation of complex, hollow networks beyond the build space of commercial tVAM devices. In many cases, it can reduce the overall dead volume used for printing by more than 50% compared to printing the whole structure at once. As proof of concept, we made a channel system recapitulating the physiological complexity of human vasculature in vitro. The system was tested with liquid flow dynamic and stent placement into stenosed segments to show that leak tight models could be created. Further, by using gelatin methacryloyl, a large-scale system out of seven single units could be created and effectively endothelialized, showing the potential of the approach for fabrication of physiological stent placement phantoms or vascular tissue models.

**Table of content figure:** 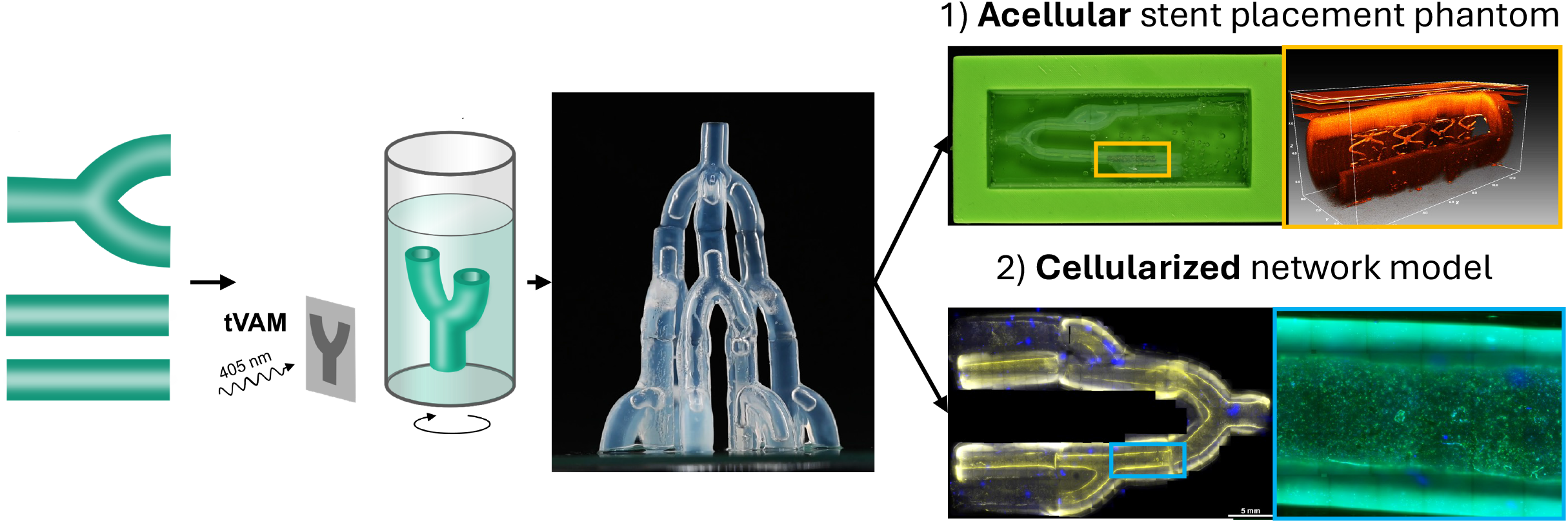

## Introduction

Additive manufacturing (AM) has found applications in many aspects of modern society ^[1]^ and specialized printers cover a vast range of different applications^[2]^. 3D printing and its derivative technologies are greatly affecting how objects are created^[3]^. Due to the commonly used layer- by-layer approach in AM, internal and interconnected structures can be realized that were difficult or outright impossible with previous technology like injection molding or milling, enabling new application fields^[4] [5] [6] [7]^. Nevertheless, there are still challenges to be overcome in the field of AM specially in upscaling and large-scale objects that, in case of light-based printing methods, require considerable amounts of resin^[8]^. Depending on the manufacturing method, complications arise like the necessity for expansive support structures or poorly draining resin from cavities^[9]^. These limitations are especially critical in terms of tissue engineering and bioprinting applications, for example the manufacturing of small to medium sized hollow tissue models. Novel manufacturing methods like volumetric printing allowing for the creation of structures without supports^[10]^ are already attempting to solve this hurdle but face process inherent limitations in terms of build size. Applications of precisely manufactured tubular tissue models are not only interesting area where hollow tubular structures are needed. This technology can be translated to soft robotics^[11] [12]^ and medical applications like in dentistry^[13]^ or surgical planning^[14] [15]^.

The field of medicine in general has greatly profited from advancements in additive manufacturing^[16]^. Many different approaches have already been explored, from hip replacements^[17]^ metal printed prostheses^[18]^, to cranial replacement options^[19] [20]^ and cellularized tissue models^[21]^ with the aspiration to eventually recreate whole functional organs^[22]^. Depending on the desired application, many different fabrication strategies have been established and explored for an eventual transition into medical trials^[23] [24]^. With more specialized 3D printing machines, highly customized and application specific constructs can be generated for restoring physiological functions in tissues and organs^[25]^.

Beyond future devices for implantation and robotics, 3D manufacturing also offers new approaches to challenges in the broader field of interventional medicine. One of such examples are vascular phantoms which are used to practice minimally invasive surgical procedures^[26] [27]^. Here, novel approaches would be needed to fabricate more life-like systems that can also replicate the mechanical properties of blood vessels to better capture the physiological situation during training of new medical personnel^[28]^. Even with state of the art manufacturing, vasculature models still pose one of the biggest challenges in biofabricated model systems. Amongst these are for example three-dimensional branched structures that allow for endothelialization to reduce platelet adhesion^[29]^. Besides better phantom models, more life-like vasculature is also integral for the development of more complex tissue models that require active perfusion for translational purposes^[30]^.

To tackle this challenge, the aforementioned process family of light-based volumetric additive manufacturing (VAM) is forming up to be at the forefront of modern tubular model systems. From the development of novel light-based printing methods in recent years, advances in tomographic multibeam AM techniques like computed axial lithography^[31]^ and holographic patterning of light fields^[32]^ initiated the emerging field of VAM as potent new technology for the rapid and freeform manufacturing of complex objects^[33]^. One subcategory of VAM, tomographic VAM (tVAM) employs a digital mirror device to project light patterns into a photo resin containing vat that rotates in a synchronized way with change of projection patterns^[33]^. Due to overlap of these low intensity projections after a full rotation, the photoreaction is initiated only in the area of maximum intensity of single projection from different angles, resulting in the generation of the object within the vat. After establishing tVAM mainly for technical resins like Polyethyleneglycol-diacrylate (PEGDA)^[32]^ it gained popularity in biofabrication and regenerative medicine research^[10]^ since it allows for the rapid creation of complex 3D-structures using photosensitive (bio-)resins in minutes or even seconds.

For medically relevant models, cytocompatible photo-crosslinkable gelatin-based hydrogel precursors like gelatine methacryloyl (GelMA)^[10]^, allyl-modified gelatine (GelAGE)^[34]^ or gelatine norbonene (GelNB)^[35]^ have been shown to be suitable and versatile matrix materials for different cell types as well as organoids^[36]^. Unlike traditional layer-by-layer bioprinting, which can be slow, tVAM fabricates the entire structure simultaneously, significantly reducing printing time and enhancing cell viability. Using this fast technique the models can be adapted, altered and exchanged by computer-aided design (CAD) modelling or between prints. Amongst other achievements, studies showed the potency of tVAM for the recreation of bifurcating and stenosed tubular structures for vascular applications^[37]^.

One major obstacle for tVAM and many other high precision manufacturing methods, is the limitation in build size. For tVAM it is especially noticeable due to exponential light absorption within the vat for the creation of larger objects not only in vertical but also in horizontal dimension^[12]^. While other VAM methods already address this issue with, for example, forwarding reservoirs for continuous fiber printing^[38]^, or light sheet printing^[39]^, tVAM relies mostly on optical corrections and intensity profiling to overcome this issue^[40][41][42]^. The currently largest manufactured objects with tVAM reach 60 mm in length and are fabricated in 100 mm diameter vats with the use of different oxygen reducing additives with potential cytotoxic effects^[43]^. Literature reported vat sizes have diameters up to 100 mm and around 250 mm height but still would necessitate additives to allow for the manufacturing of large scale objects based on high resolution resins^[43]^. With larger vats, the total amount of required photoresin is also increasing and to ensure maximum print fidelity is ideally not reused after a finished print^[44]^. This results in large dead volumes and discarding of expensive printing materials, which further amplifies the costs of this already expensive method for smaller laboratories.

To overcome the current limitations of tVAM and harness the flexibility of the tVAM method for large scale hollow structures, we established a manufacturing workflow that allows the generation of complex hollow branched geometries by utilizing customized small building blocks (modules) (Fig. 1). These can be easily attached to each other by reinforcing the connection with photoreactive materials to create structures beyond the build volume limitation in centimeter scale. We tested the system with non-cellularized and cellularized approaches for vascular model systems, requiring leak tight branched tubular structures with physiologically relevant inner diameters. As a demonstration for small diameter vasculature networks, assemblies were created that feature narrowing down diameters in accordance with Murray’s law^[45]^. As a non-cellularized model, a tubular assembly with stenosis has been created to serve as a stent placement phantom. Then, as a cellularized model, an assembly of modules has been cultivated by seeding the building blocks with human endothelial cells before assembly. By that we could demonstrate that the system is capable of creating large endothelialized structures with high cell density on the lumen of the model, enhancing seeding efficiency of biofabricated in vitro systems and future bioassembly approaches.

**Figure 1:**
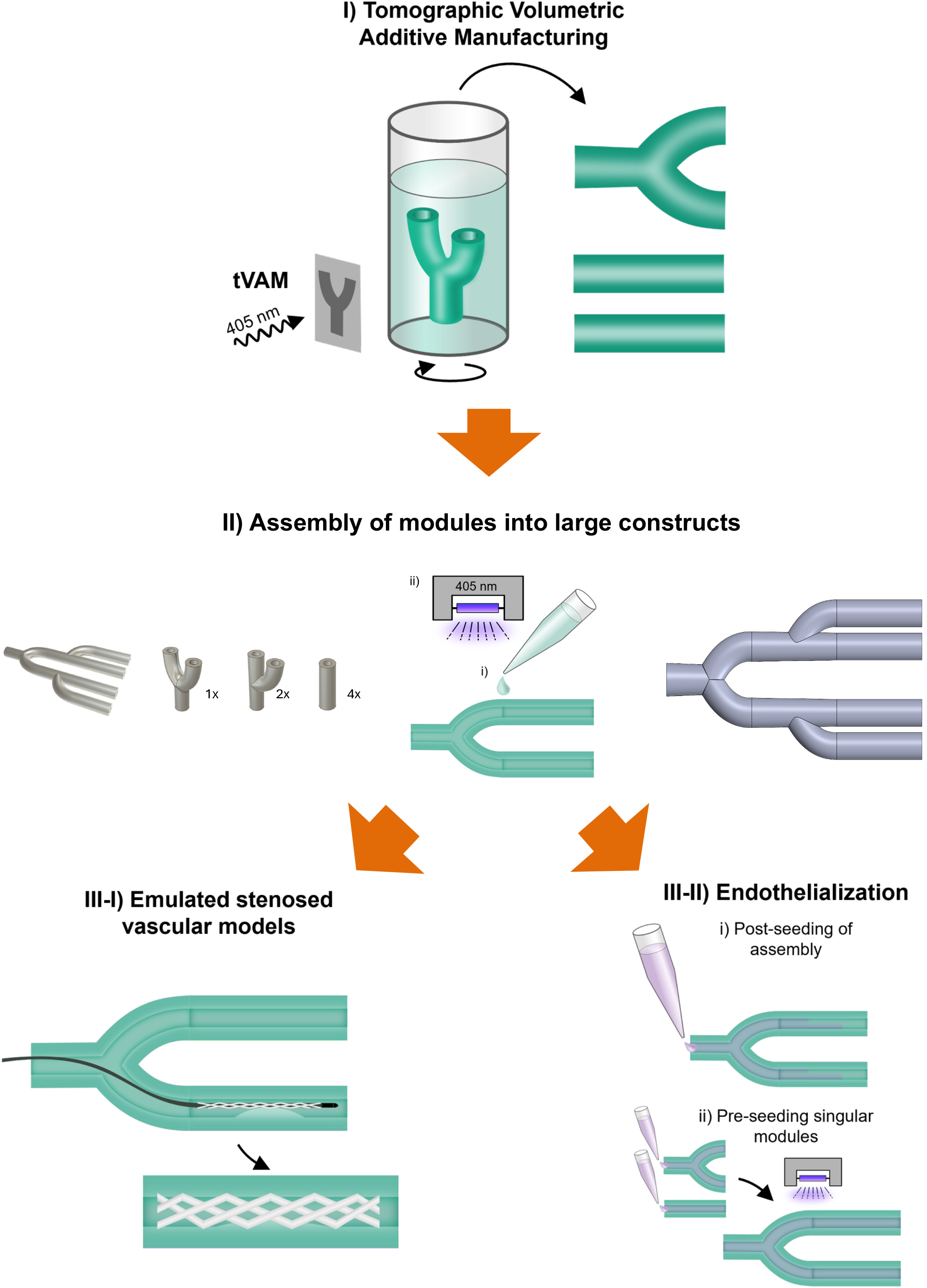
Overview of modular building block system created by tVAM. **I)** An assortment of different modules was created with different specifications that can be combined seamlessly. **II)** Assembly of single modules is done via application of a small amount of photoresin on the connection sites before crosslinking. Assembled modules are leak tight and suitable for perfusion of liquids. To achieve large perfusable model systems, multiple modules of different geometries can be combined into complex hollow structures. **III-I)** Assemblies out of PEGDA are suitable to emulate stenosed vascular networks for the placement of balloon-catheter deployed stents. **III-II)** Assemblies out of GelMA allow for pre-endothelialization with different endothelial cell types from mouse and human. By pre-seeding single modules, a high degree of endothelial covered luminal area can be achieved compared to a post-seeding of a whole assembly. Further, perfusion of modules alters gene expression of HUVECS cultured within single modules, underlining the future potential of this workflow for both, acellular and cellular model systems of vasculature.

## Results and Discussion

To verify the initial hypothesis of this approach, a set of tubular modules with different complexity levels have been designed using CAD software to assess the precision and reproducibility of tVAM. As tVAM is based on the projection of rotating images into a photosensitive material (Fig 2A), the assumption was that slight variations in material or centering of the vial within the printer could affect the reproducibility of single units which would be detrimental to the manufacturing approach.

**Figure 2:**
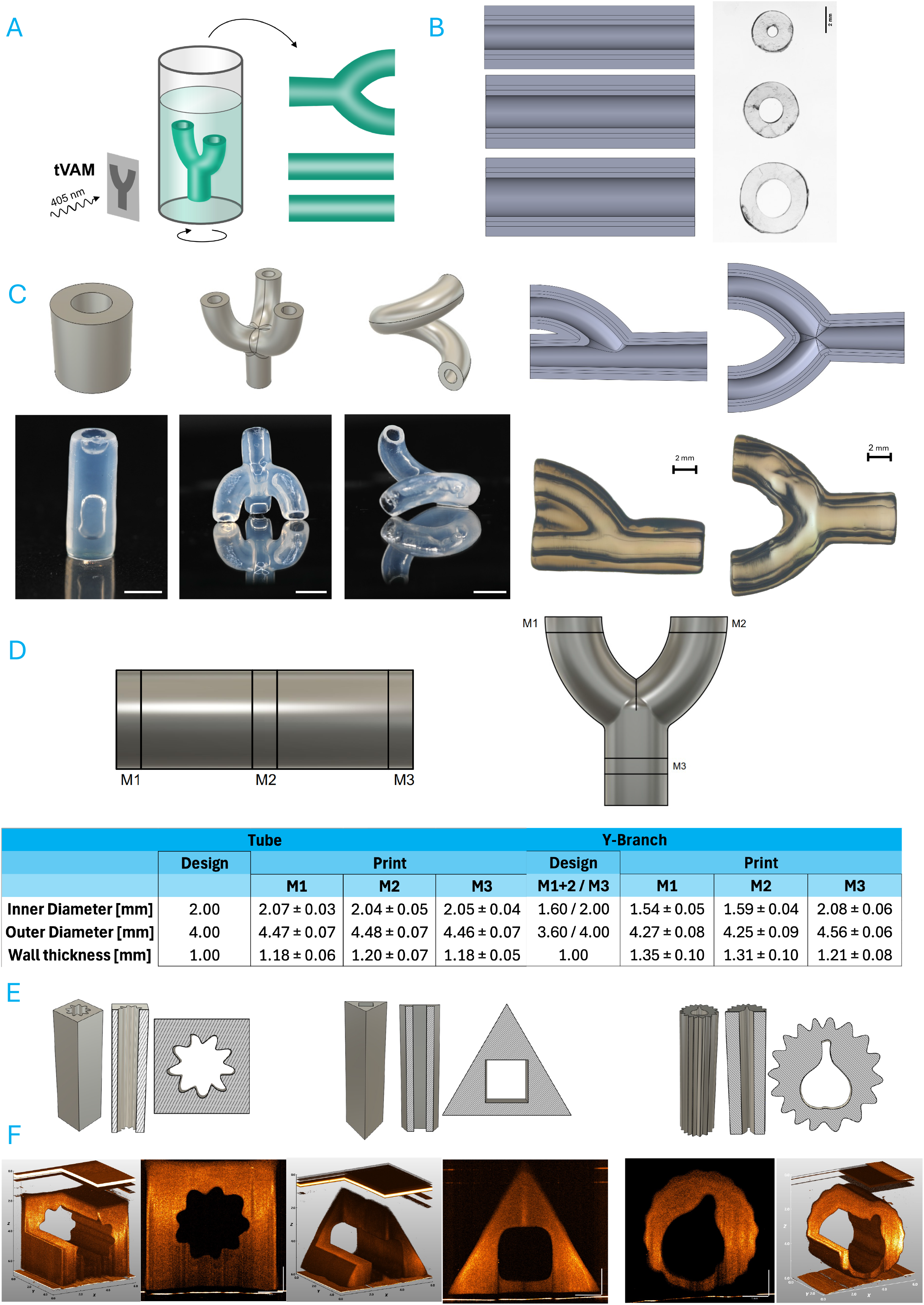
Manufacturing precision of modular building blocks created with tVAM. A) Schematic representation of tVAM fabrication of modules. B) Design and manufacturing of varying inner diameter tubular structures. CAD designs of 1, 2 and 3 mm inner diameter structures with 1 mm wall width compared to printed construct cross sections (SB = 2 mm). C) Design and manufacturing of complex tubular modules. CAD designs of tube, claw, macaroni, h- and Y-Branch and the achieved printed structure (SB = 2 mm). D) Evaluation of print accuracy for different tubular modules. M1-3 indicates measurement areas (N = 5, n = 8) E) different possible CAD designs for single modular hollow modules. F) verification of printability of CAD models via OCT imaging of a side view as well as a luminal view (SB = 1 mm).

As a material, polyethylene glycol diacrylate (PEGDA), a commonly used synthetic resin for tVAM, with the addition of cellulose nanofibrils (CNF) as rheological modifier has been chosen for an initial proof of concept study. A study with different amounts of PEGDA and the addition of 0.24 % wt/wt CNF was carried out to evaluate if the addition of CNF affects the material properties and diminishes printing quality. The polymerization characteristics of PEGDA concentrations (2.5, 5, 7.5, 10 and 12.5 wt%) was evaluated via photorheological measurements. All concentrations with the addition of CNF showed polymerization upon exposure to 405 nm wavelengths with LAP 0.05 % as a photoinitiator, with the exception of 2.5 wt% (SFig. 1 A). An increase in Young’s Modulus was measured increasing with polymer concentration, showing an exponential increase in stiffness relative to the increasing measured PEGDA concentration (5 wt%: 3.7 ± 0.7 kPa; 7.5 wt%: 22.3 ± 4.4 kPa; 10 wt%: 65.4 ± 12.0 kPa; 12.5 wt%: 173.6 ± 32.7 kPa) (SFig. 1 B). An increasing turbidity of the investigated material formulations could be observed relative to the amount of used PEGDA, where 5 % produced the clearest gel structures (SFig. 1 C). Since maintaining a high precision of the luminal space of each module was a priority for the intended application, and a turbid material could affect light propagation throughout the materializing structure, the 5 wt% PEGDA concentration has been chosen as material to use for further evaluations.

To put the results into perspective, one has to consider that photopolymerization by tVAM allows for precise spatial control and fast printing times in the range of seconds to minutes^[10]^. By variation of the photopolymer content, the mechanical and optical properties of the hydrogel precursor solution and the resolution of the resulting hydrogel can be tailored to fit the intended application (SFig. 1).

PEGDA hydrogel-precursors solutions^[46]^ modified with CNF^[47]^ as a rheological and structural enhancer, shown in other works employing a similar approach^[48][47]^, are ideal for vascular models, which provides a cost-efficient platform for various biomedical applications The high transparency of the PEGDA leads to a rapid and controllable gelation of the polymer resin through photopolymerization which makes it the perfect resin for tVAM. We chose PEGDA 700 for the model as a well-researched, readily available and synthetic polymer with non-cytotoxic properties as a cheap alternative to natural and other biocompatible polymers like collagen and gelatin derivatives because of its high water-solubility and well defined structure, high batch-to- batch consistency and molecular weight. PEGDA has been used as a low cost resin for tVAM before with the addition of alginate as a rheological modifier^[49]^. A thickener is necessary since low molecular PEGDA since the intrinsic viscosity is too low to keep polymerizing structures suspended in their location during tVAM printing^[49]^. The nanocellulose content adjusts for viscosity and prevents objects from sedimentation and counteracts turbulence and diffusion in the resin in print^[47][49]^. To guarantee a decent printing quality, transparency of the resin is highly relevant. Optical modifiers can be added to adjust for inclusion of other materials or particles that could affect transparency of the materials, especially when used in conjunction with cells^[50]^. Even though the used CNF are within nanometer scale, an elevated amount within the photoresin blend will be noticeable as increasing opacity. In a test with increasing CNF amount in PEGDA ranging from 0.08 % wt/wt to 1.2 % wt/wt it was evident that concentrations above 0.3 % wt/wt CNF decreasing transparency of the resin (SFig. 2A), affecting also the print fidelity and surface structure (SFig. 2B). For optimal results a concentration of 0.24 wt% wt/wt was chosen, which resulted in the overall best results in surface quality, transparency and print fidelity and a rheological modification comparable to the addition of 1 % wt/wt xanthan gum (SFig. 2C).

With the material for manufacturing chosen, a study on different diameters of cylinders has been conducted to ascertain a precise manufacturing of different inner diameters. The screening of luminal diameters from 1 to 3 mm with a wall thickness of 1 mm in each condition revealed minimal deviation from the CAD design in all conditions (Fig 2B and SFig. 3A). Each tube was cut and measured eight times at three different places (M1-3) (Figure 2D). Tubes with a theoretical inner and outer diameter of 1 mm showed an actual inner diameter of 1.02 ± 0.02 mm, an outer diameter of 3.20 ± 0.04 mm and a wall thickness of 1.09 ± 0.04 mm. For the tubes with a designed inner diameter of 2 mm and an outer diameter of 4 mm, the measured inner diameter was 2.05 ± 0.04 mm, the outer diameter 4.47 ± 0.07 mm and the wall thickness 1.19 ± 0.06 mm. The largest modules featured an inner diameter of 3 mm and an outer diameter of 5 mm. The recorded measurements were for the inner diameter 3.02 ± 0.05 mm, outer diameter of 5.66 ± 0.10 mm and a wall thickness of 1.29 ± 0.08 mm. All measurements correspond closely with the CAD model, especially the inner diameter.

**Figure 3:**
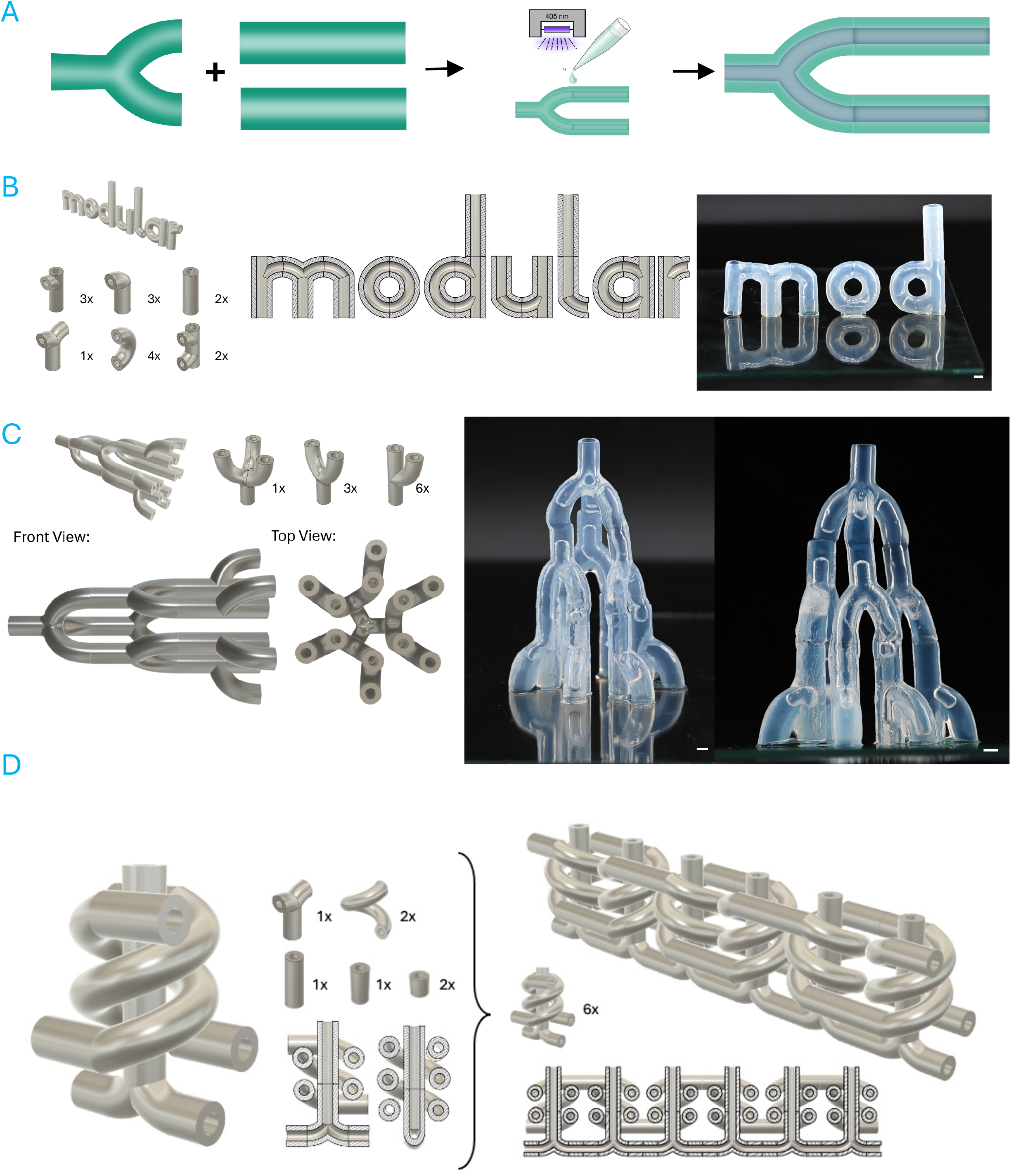
Design space of hollow network structures fabricated by modular tVAM. A) Schematic overview of the assembly process of multiple modules. B) Potential of customizing single modules to create a wide variety of assembled models. Example shows the writing ‘modular’ recreated with single modules (SB = 2 mm) C) CAD model and assembled three-dimensional assembly. With the three branch claw module as center piece, a ten module assembly has been created which utilizes the 3D space than conventional perfusion channel systems (SB = 2 mm) D) Complex, multi-module assembly of a three-way perfusable system. The modularity of design possible via tVAM allows for concepts of module systems that could be connected to create large perfusion models.

Next, the freeform manufacturing of tVAM was utilized to create not only simple cylindrical tubes, but also Y-branched modules with narrowing branches in accordance with Murray’s law for vascular systems (Fig 2C). To further tap into the potential of tVAM, volume structures beyond simple cylinders that better utilize the geometrical freedom of X-, Y- and Z-dimensions have been tested. For this reason, the shape of a three-way-branched module (claw), a helically coiled module (macaroni), as well as two different kinds of bifurcations, one with a bypass (h-Module) and one naturally splitting (Y-Module) have been designed. All the designs were manufacturable via tVAM and the produced results were comparable to the CAD design (Fig 2C). To verify adherence to the CAD design after manufacturing, regular cylindrical modules as well as narrowing branched modules have been evaluated. Similar to the measurements carried out for Fig 2B, an increase of the outer mantle area has been recorded for all measured modules, whereas the deviation for the inner diameter was below 5% compared to the CAD design (Fig 2D). The Y-Module designed with narrowing, according to Murray’s law, featured inner diameters of 1.60 mm at the branched section, whereas the main channel featured an inner diameter of 2.00 mm. The inner diameter of the smaller channels (M1 + M2) with 1.54 ± 0.05 mm and 1.59 ± 0.04 mm closely replicate the designed 1.60 mm. The main channel (M3) with 2.00 measured 2.08 ± 0.06 mm close to the dimension of the model as well. The outer diameter with 4.27 ± 0.08 mm and 4.25 ± 0.09 mm for the branch channels and 4.56 ± 0.06 mm for the main channel shows slightly more deviation from the original model.

While for most applications in biomedical fields, a tube-like structure is required, different shaped hollow channels might be beneficial for other applications. To show the feasibility, a selection of different shape combinations for hollow modules has been evaluated and verified via optical coherence tomography (OCT). Two aspects were evaluated, outer and inner geometrical shape. With an outer shape of a square or triangle, parallel channel systems could be easily stacked on top or next to each other, while a toothed shape might be interesting for kinetic models that rotate in conjunction with each other (Fig 2E). The luminal shape is highly dependent on the manufacturing resolution of the method. Here, tVAM excels with replicating shapes with features of a cogwheel or a Franconian wine bottle, indicating that virtually any shape might be recreated to alter flow kinetics within the module, or the whole assembly (Fig 2E,F).

The advancement of tVAM technology enabled it to become a precision manufacturing technique, approaching resolutions as small as 50 µm^[12][36]^. Novel mapping and AI-based calculation models further enable precise control over the resulting print, even enabling tracking of objects within the vat^[51]^. Reported resolution limits range down to 31 µm^[52]^ whereas new technological advancements of the technology managed to reach up to 12 µm of resolution^[53]^. It is, nevertheless, highly dependent on the employed materials and overall transparency of the resins, directly affecting the outcome and quality of the print^[50]^. The addition of CNF into PEGDA did not negatively impact the resolution of the printed modules, which allowed for complex geometries to be created via tVAM. As previous studies of low molecular weight PEGDA have already showed, a rheological modifier is necessary for processing with tVAM^[49]^, CNF is a viable modifier for this approach. Regarding accuracy to the model, tVAM achieves deviations of below 1% in print deviation^[54][43]^. The reported measurements of modular units are in the range of 1-8 % deviation, depending on the model. This is a moderate increase compared to highly optimized systems and could be caused by the inclusion of the nanocellulose. Deviation compared to standard non-corrected tVAM, which can have up to 50% deviation, this small increase compared to the cutting-edge standard can be considered as small^[55]^.

While the created modules are not able to recapitulate capillary networks, which would require channels of sub 50 µm diameter and is the main focus of modular microfluidic systems^[56]^, it does open the field to larger scale vessel systems. Since tubular model systems are usually used to recapitulate vascular networks, the presented study is highlighting possible future applications of a modular building block system for small diameter blood vessels^[57][58]^ and synthetic blood vessel systems^[59]^. The impact customized vascular systems can have on different medical fields, especially for cardiovascular applications has been positively evaluated^[60]^. With the use of volumetric printing technology, previous obstacles like complex geometries or strong layer-by-layer artifacting or the need for support structures of other printing techniques can be circumvented. One clear advantage is shown by the seamless integration of vessel narrowing in bifurcations described by Murray’s Law^[61]^ and complex free-standing geometries like the presented macaroni module. 3D printing enabled new possibilities of geometric shapes with interlocking or kinetic properties for movements^[62][63]^. Specialized workflows and geometries also facilitate the generation of interlocking modules that allow kinematic movements of the assembly^[63]^. Early on, tVAM also was shown to be able to manufacture such shapes with the example of a ball-cage valve where an incorporated sphere acted as an actuator to regulate flow^[10]^. Similar strategies would also be transferable to single modules with specialized functions, as for example regulate flow speed or directionality. The latter was also shown in previously published work in the form of geometries that acted as functional venous valves^[37]^.

In summary, a cost-effective material combination in the form of PEGDA 700 with 0.24% wt/wt CNF was found that allows for precise manufacturing of tubular modules of different geometrical complexity. The inner diameters of the printed constructs were manufactured with high precision in relation to the dimensions of the CAD model, both for cylindrical modules as well as the Y-bifurcation module. These initial series of experiments indicate a high likelihood of compatibility between single modules when manufactured with tVAM.

With single module complexity explored, a connection between the single elements was investigated to elucidate if a modular building-block-like approach to create larger tubular models would be possible with tVAM printed modules.

We observed that when two modules are put close to each other, they tend to link together on their own. This preliminary weak attachment is used for adjustments of alignment before fixing the layers using glue (Fig. 3A). For gluing a thin layer of the same photocurable resin is applied to the connection site (SFig. 4A, C). If the print displayed uneven edges towards the in or outlet, a specialized cutting mold created with digital light processing (DLP) was employed to trim the edges for a flush fit of two modules (SFig. 4B). Further, to aid in alignment, specialized molds were tested to place the modules in the intended assembly and glue them without the risk of displacement during curing (SFig. 4A). Stiff synthetic DLP resins were also tested and resulted in a stable connection of modules but due to the difference in stiffness, between the cured resin and the hydrogel, same material adhesion was preferred.

**Figure 4:**
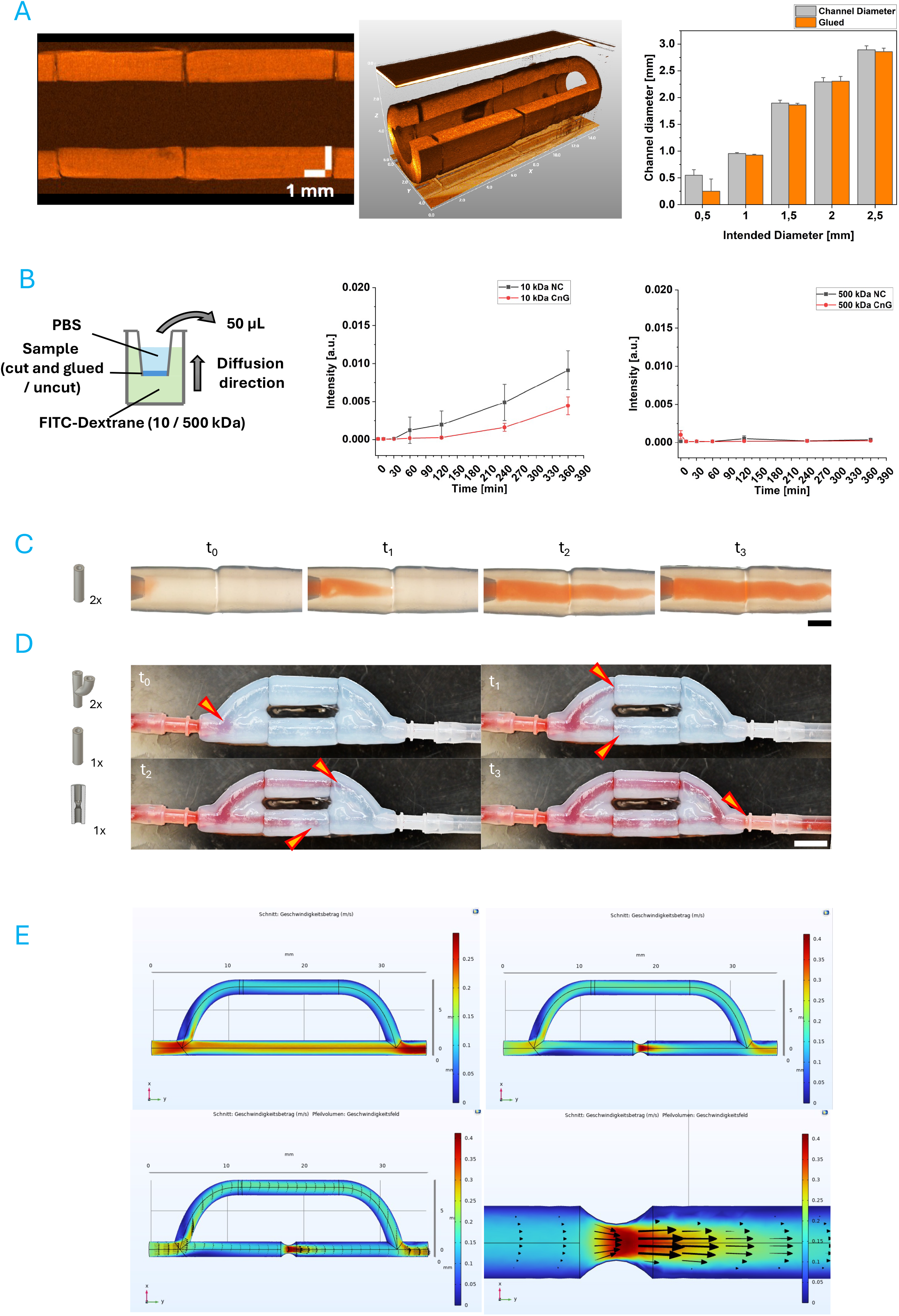
Suitability of the modular system for perfusion and stenosis models. A) Representative OCT image of multiple short modules combined for inner diameter measurements after assembly. Both a vertical section and a tilted overview are shown. Bar graph shows statistical analysis of changes of inner diameter after assembly dependent on initial fabrication diameter. B) Simplified diffusion experiment of FITC-Dextran through cut and uncut hydrogel disks fabricated with volumetric printing. The schematic illustrates experimental setup for diffusion of 10 kDa (first graph) and 500 kDa (second graph) through uncut (NC) and cut and glued (CnG) hydrogel discs to mimic the attachment of tubular modules. C) Time points (0 = 0 sec, 1 = 5 sec, 2 = 10 sec, 3 = 15 sec) of perfusion experiment of colored water through an assembled multi-module tube structure (SB = 2.5 mm) D) Time points (same as B) of perfusion experiment of colored water through a stenosis (indicated by blue dotted line) and bypass multi-module assembly (SB = 10 mm) E) Computational fluid dynamics of the assembly in D, with and without stenosis (top graphs) as well as flow direction in the stenosed model (bottom graphs). Scale in m/s.

Regarding the assembly of more complex modules, the feasibility of different models was tested on a general level initially. The aim was to create the word ‘modular’ with several single modules and continuous lumen. All designed assemblies would not be possible to be created with the stock vial size available on the commercial tVAM printer used in this study, either due to size limitations or refraction artifacts at the outer interface of resin and glass. As depicted in Fig 3B, the CAD designs all feature a continuous lumen to theoretically allow perfusion with liquid, with the exception of the O. Specialized modules were designed to create the geometrical shapes of letters, with ‘r’ shaped modules present in the ‘m’, ‘u’ and ‘r’ letters, a module combination to create an angled 90° turn in the lower ‘l’ as well as modules with one-sided double bifurcations present in the ‘d’ and ‘a’ (Fig 3B). The first three letters were also printed and assembled as seen in the second part of Fig 3B.

A significant advantage of additive manufacturing is the effective use of all three dimensions for designs. To harness this potential, the previously presented ‘claw’ module was used as center piece for a three dimensional assembly of modules. The design as well as the final assembly of all modules is shown in Figure 3C. The assembled pieces defy gravity and maintain their shape in the vertical direction.

Conceptually, the presented modular system can also be used to create intricate shapes which would be difficult to recreate with any other manufacturing technique. Figure 3D shows a hypothetical design with three separate perfusion loops that can be assembled in one unit but can also be connected to longer complex assemblies.

Assembly of modules and gluing technique worked as shown in a similar publication using modified GelMA as a matrix material for building blocks^[64]^. Compared to that publication, the geometrical shape of tubes with little attachment area, the attachment was expected to be more challenging than the previously shown cubic modules^[64]^. The presented results of this study illustrate that the method works just as well with tubular modules. For the assembly of multiple modules, an alignment tool was printed using DLP technology to easily match to segments and apply the resin to join them (SFig. 4A).

Volumetric printing excels with its ability to create complex shapes in very short printing times^[10]^. The presented geometries illustrate the adaptability of modules with simple CAD designs that also seamlessly fit together to build complex assemblies with perfusable lumen. Moreover, the possibilities are not restricted to purely biological medicinal designs but also of technical nature as illustrated by Figure 2E and 3D. The simple concept of joining modules and fixing them with photoresin enables the operating person to quickly assemble a desired geometry after printing the required modules (Fig. 3C). While most perfusion applications are created with a planar 2D design, 3D printing enables the quick generation of multi-level channel systems that can be situated above, below or even around the main channel network^[65]^. Figure 3C highlights the possibility with the presented modular system to create channel networks that feature more complex three-dimensional architectures. Such assemblies could be used as hydraulic grippers with redesign of the single modules to serve as pressure driven actuators^[66][67]^. Due to the increase in size possible with this modular approach, also larger systems with highly precise geometries for intricate articulation could be realized. In this study, only larger channel modules were tested but tVAM could be used to create intricate interconnected small channel modules similar to the designs presented by Gregoryan et al.^[65]^. The final assembly was done outside of an immersive medium, resulting in selective dehydration of the PEGDA hydrogel that displaced some of the modules over time. For larger assemblies like this, a suitable immersion within an aqueous solution could be beneficial to maintain the original geometry for as long as possible. It would also aid in maintaining a stable positioning of the assembled units while positioning the next ones. If such a complex structure were to be attempted in the future, submerging it in a support bath consisting of for example xanthan gum would likely facilitate the assembly while enabling photo-crosslinking and being overall cytocompatible^[68]^. Nevertheless, the presented assembly consisting of 10 single modules highlights that the system can also be used without any further supporting systems.

By utilizing the approach of printing single modules instead of the whole structure at once, the printing time naturally increases since each module requires its own printing run instead of one run for the whole structure. This allows for better resolution of the overall assembly and avoiding the complications of light intensity falloff in large vials, further enhanced by scattering and refraction at the glass/resin interface of the vat^[43]^. Further, printing with smaller vials and effectively using the build volume also reduces the overall material consumption required for one large structure. Assuming resin is only used for one print (due to the chain-growth radical polymerization of most methacrylated resins that affect print fidelity when used more than once^[44]^, depending on the structure printed, a certain amount of dead volume will be present that would be discarded. By assuming the standard cylindrical vat geometry, due to diffraction at the interface, the maximum printable volume accounts to around 62.5 % of the total volume of the vat. When calculated for the size of vat required for the presented structure in Fig. 3C, the required vat size would be 60.8 mm and includes plenty of unused volume due to the branching structure that is widening towards one end. By using smaller vats (10, 15 and 20 mm diameter) for each module, the build volume is used more effectively and overall unused volume is reduced, resulting in a reduction of 70.9 % required resin if the above assumed 62.5 % of effective build volume is considered. This significantly reduces the required costs for printing with expensive or hard to manufacture photoresins and empowered the technology of tVAM even further for novel experimental materials and economical use of the method. These aspects are especially relevant in case of bioprinting applications where bioresins containing cells are used. While only conceptually illustrated in Figure 3D, the potential of this tVAM generated modular system can be employed to generate complex interconnected structures that have not been able to rapidly be fabricated before. Further optimization in perfect alignment of the non-planar structures would be required to ensure a proper interconnectivity of the perfusion lumen which could be realized with spacers or assist devices printed with the negative shape of the module (as shown in SFig. 4). The presented geometries highlight that this proof of concept system can be realized and transferred to several different applications, both technical as well as medicinal. Regarding technical applications, recreating physiological structures is also a core aspect of the field of soft robotics that attempt to mimic natural shapes as actuators^[69]^. Fluid based kinematics are especially interesting due to their rigidness, ease of fabrication and high gripping force^[70][71]^. A new class of endoskeletal soft robots inspired by physiological structures also rely on a precise geometric recreation of hollow network structures^[11]^. The concept presented here, especially the ‘claw’ module which mimics a common three-feature gripper, could be transferred in future applications to an actuator assembly with highly customizable gripper designs that are either hydrodynamically or activated or via precision swelling of the hydrogel component as demonstrated before^[11]^.

The above presented concepts illustrate that the module system offers high customizability of single elements which enables creation of tubular assemblies of varying complexity, both in 2D and 3D. The future implications of such a model range from simple branched tubular networks up to complex vertically and horizontally scalable perfusion network systems. The quality of connection between single modules was investigated by OCT measurements of the connecting site between two modules. Linking two or more modules together was found to be a straightforward process involving a gluing step to attach two or more modules as shown in Figure 3. To further confirm that the luminal area is not affected by the application of additional photoresin, OCT images of the connection site have been recorded to monitor if resin is entering the luminal area, affecting the overall diameter (Fig. 4A). Modules with varying inner diameters (0.5, 1, 1.5, 2, 2.5 mm) have been attached to each other and measured via OCT in a longitudinal cut and as a full 3D scan displayed from a side view. (Fig. 4A). No significant changes of diameter were recorded for diameters of 1 mm and above, but for the smallest diameter of 0.5 mm the channel diameter was reduced to about half after gluing (Fig. 4A).

Overall, the way of assembly presented is fully compatible with hollow tubular modules without affecting their luminal diameter. The reduction measured in the smallest tested diameter is likely due to small amounts of resin entering the lumen and accumulating due to the smaller luminal area and by that higher surface area where surface tension dominates and promotes droplet formation, resulting in reduced inner diameters up to clogging. In larger diameters, this leaking resin redistributes itself more evenly across the larger luminal surface, likely also creating a smoother surface on the inside. Since this phenomenon was not present in all investigated samples, as shown by the standard deviation, it might also be affected by the operator’s consistency of applying an exact amount of resin onto the gluing site of the module. Further structured investigations for the suitability of this system with smaller diameters are necessary in the future to ascertain a use case for hollow structures also at 0.5 mm and below.

To assess the leak tightness of assembled modules, a custom diffusion assay using FITC-Dextran was performed. A DLP-printed well plate insert was used that fits small material discs to separate the well reservoir from the underlying well (Fig. 4B). To simulate the leak tightness of assembled tubular modules, the discs were cut in the center and reassembled in the same fashion as modules would be attached to each other. Discs in uncut state and with a glued cut were assayed with several molecular weights of dextran (10, 500 kDa) dissolved in liquid, which was inserted into the well. Measurements of the liquid within the upper reservoir were carried out to quantify the amount of FITC-Dextran diffused through the hydrogel disk. Both molecular weights of FITC-Dextran were retained by the investigated hydrogel formulations, both in cut and uncut condition, illustrated by the near zero value of the fluorescence quantification. Cutting and regluing resulted in a slower increase in fluorescence quantity showing no increase in diffusion through damaging the hydrogel. Further, it was investigated whether the results of the simplified study could be translated to the actual tubular system. For this purpose, assembled modules were pre-filled with water and subsequently perfused with red food dye. An assembly of two strands was tested, and pictures were taken at different time points (0, 5, 10 and 15 sec) of the experiment after perfusion to visualize the progression of the perfused dye (Fig. 4C). The assembled construct showed no leaking during or after the experiment.

Diffusion of molecules through hydrogels is widely studied and is dictated predominantly by hydrogel composition and a number of factors like degree of modification and molecular dispersity among others^[72]^. The utilized setup is based on an already established system called Franz diffusion cell and is an established method in pharmacology^[73]^ and overall scientific diffusion and release studies of hydrogels^[74]^. To assess diffusion out of hydrogel-based systems, utilizing water-soluble molecules linked to fluorescent dyes is a common practice^[72]^. FITC-Dextranes found wide applicability for this purpose due to their inert character and their breadth of available molecular weights, specifically relevant for release studies^[75]^. The employed PEGDA itself formed a dense hydrogel network that prevented the perfusion of the investigated dextrane weights, making it an ideal choice for most non-biological use cases of the modular system where liquid perfusion would be required^[76]^. For future applications, different formulations of PEGDA could also enable the selective diffusion of molecules or act as compound carrier to release solutes into the luminal cavity via diffusion, further enhancing the potential applications of the presented system^[76]^.

After the baseline evaluation of leak-tightness of hydrogel formulation and assembled building blocks, a more functional model with a stenosed module was created to show a potential field of application for the modular system. For this purpose, a stenosis was added to the middle of a straight tube in its CAD design with dimensions of 0.5 mm in diameter (effectively reducing lumen of the tube by 1 mm), a width of 3 mm and was radially symmetrical (Fig. 4D). To verify the effect of the stenosed module, a bypass assembly has been designed and assembled (Fig. 4D). A perfusion setup with thorn barbs and silicone tubing has been attached on both ends of the assembly and perfused via a dye loaded syringe. During the perfusion, pictures at multiple time points were recorded to monitor the dye flow. Shortly after beginning of the experiment, the liquid flow was obstructed by the stenosed segment in the main vessel (Fig. 4D t1). Due to the congested flow, the perfused liquid was redirected through the longer bypass as seen in t2 and t3 and merged again after the stenosed module (Fig. 4D). These findings were confirmed by flow simulations that show higher average flow rates in the bypass than after the stenosis module (0.2 - 0.25 m/s compared to 0.1 - 0.15 m/s) (Fig. 4E).

The findings of this experimental setup are in accordance with comparable bypass models in literature^[77] [78] [79] [80]^. This proof of concept verifies that stenosed vessel systems can be recreated by an assembly that corresponds with CFD simulations. With the customizability possible with tVAM different stenosis morphologies could be created in future adaptations of the system to change the flow profile within the assembly^[81]^.

Overall, the use of 3D printing for stenosis models has been explored before and found that with the correct printing parameters, calculated values can be closely matched^[82]^. This aspect could also be investigated in future studies on translatability of this system towards patient specific recreation of stenosed vessels^[83]^. Ascertaining a predictable flow and luminal pressure would also be relevant for a potential application in soft robotics since differences in applied pressure are responsible for the actuation of the construct, further paving the way for a technical application of this system^[67][84]^.

The experimental results of perfusability of modular assemblies highlight the suitability of the presented method for perfusion networks, which align with comparable published results^[64]^. Not only could the single segments be attached together in a straightforward manner, but they also created a leak tight connection and could be used to verify flow simulations of a stenosed network system.

So far, several proof of concept experiments have been presented to illustrate the feasibility of a modular based approach for tubular assemblies created with tVAM. In the next step, potential future applications of the systems will be showcased. The focus will be on vascular-like systems for clinical use. Two use cases have been chosen to showcase the adaptability of the system: A technical application without cells that uses PEGDA as module material, and a cell-based application with GelMA as module material.

Beginning with the technical model, a stent placement model with multiple bifurcations was created. As a starting point, a vascular tree assembly has been designed to capture the complexity and length of a minimally invasive catheter intervention training model, similar to a vascular phantom. The intention was to recreate a phantom with higher complexity that can serve as a training dummy for medical students and surgeons that does not rely on straight silicone tubing and offers the option to place stents within bifurcated branches. Figure 5A shows the schematic CAD template and the tVAM printed result with all modules glued together. The total length was approx. 80 mm and included several bifurcations to emulate complex vasculature. As highlighted by the orange boxes, regardless of the complexity of the used modules the junction sides displayed a high level of fitment and precision in the gluing process. To further emulate a phantom model, a second two-branched tube system was designed (Fig. 5B). This assembly was embedded in silicone to make handling easier while providing visual reference of where the catheter is currently positioned within the system. A balloon-catheter with a loaded stent has been introduced in both arms by an untrained person to demonstrate ease of use of the model as well as its feasibility as a stent placement phantom (SVid. 1). Due to the branched nature of the model, it allows multiple pathways to navigate the catheter towards the stenosis (Fig. 5B).

**Figure 5:**
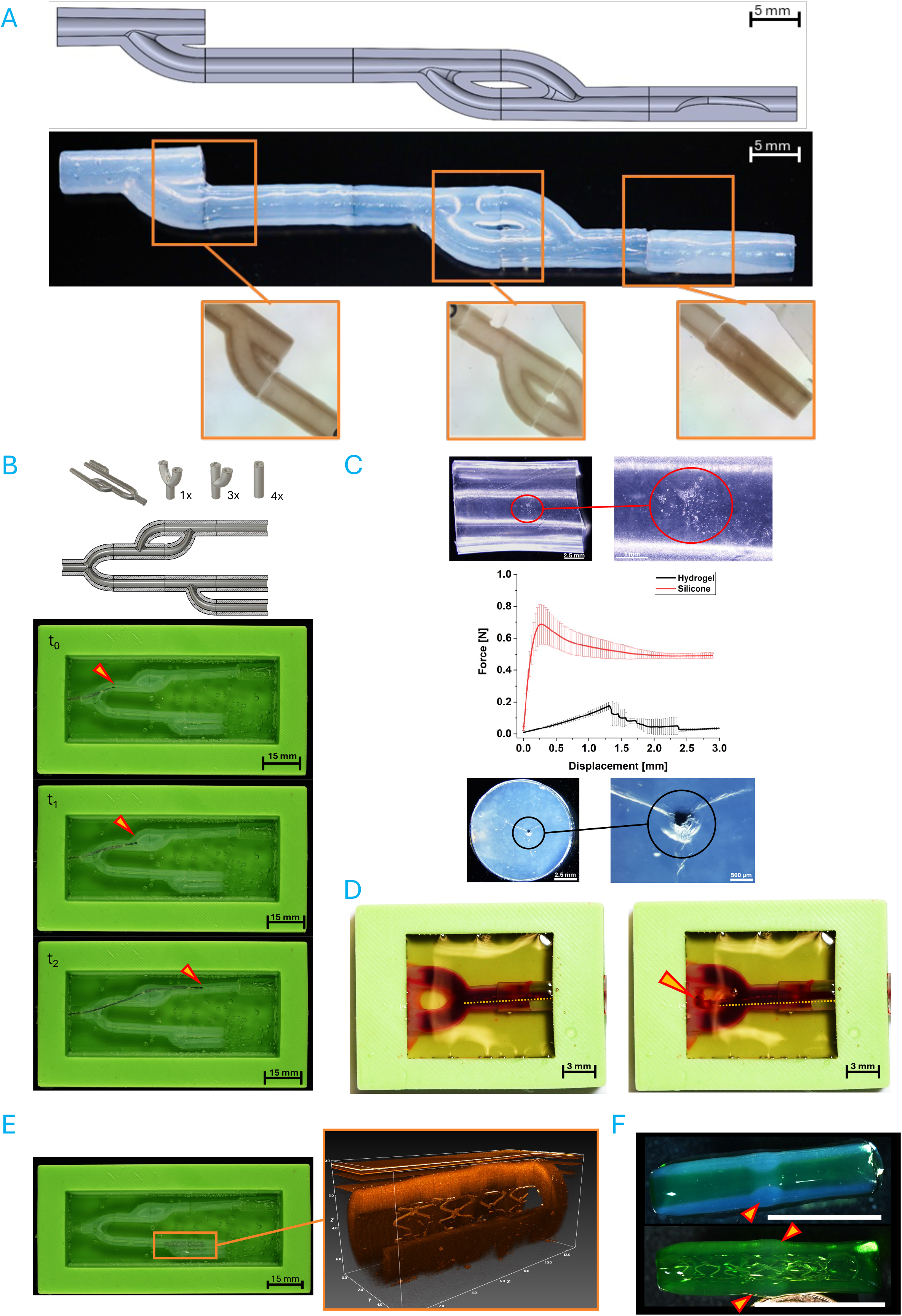
Multi-module system for the use as stenosed vascular phantom. A) CAD design and print of a large, multi-module assembly with a stenosed module at the end. Orange squares indicate where the zoomed in images were taken from the assembly. B) Overview of assembly and use as phantom for catheter navigation. The module assembly out of PEGDA has been encased in clear silicone and subsequently in green silicone for handling. An opening on the left side allowed entry of a commercially available balloon catheter. t0-t2 shows the insertion of the catheter into the system. C) Catheter force required to penetrate conventionally used silicone tubing for phantom models and hydrogel formulation used for the manufacturing of tubular modules. Red shows data and representative silicone sample after the test (upper section) black shows data and representative hydrogel sample after the test (lower section) D) Demonstration of catheter induced penetration of modular module. Allura red stained PBS in the lumen is exiting the module after the catheter (indicated with orange dotted line) pierces through the module wall (indicated by arrow). E) The lower branch of the module assembly was chosen as the site of stent placement. A commercially available stainless-steel stent could be opened into the lower branch of the assembly and verified via OCT imaging (orange square). F) Confirmation of stenosis widening after expansion of a stent within a stenosis module (stenosed area denoted with arrow). The second image shows how the stent has moved the stenosis out of the lumen and is now protruding out of the outer mantle of the module (arrow). Green food dye has been used as contrast enhancement (SB = 1C.25 mm).

Further, the penetration resistance of the modular system was compared against commonly used silicone tubing for custom made vascular phantoms. A balloon catheter with a stent was mounted onto a uniaxial testing machine and moved towards the samples while recording the force exhibited by the contact. As seen in Figure 5C, the PEGDA hydrogel was cleanly punctured, which is confirmed by the decrease of force measured after a displacement of 1.25 mm. The silicone sample on the other hand could not be penetrated by the catheter, instead the catheter bent and moved out of the way, scrubbing the surface of the silicone (Fig. 5C). This behavior is indicated by a steep increase in force that results from the compression of the catheter until it bends away shortly after 0.4 mm and then pushes and moves away from the sample, which is recorded as a force plateau of around 0.5 N (Fig. 5C). To further illustrate the damage and penetration capabilities of the modular system for closer physiological phantom models, a Y-module has been closed off at the split ends, filled with a red dye and submerged in a silicone holder (Fig. 5D, SVid. 3). By inserting a balloon catheter from the single end into the lumen and pushing it with some force into the split of the module, the catheter breached the wall of the module, releasing red dye into the surrounding medium, indicating damage to the module during the insertion process (Fig. 5D, SVid. 4).

Finally, the deployment of a stent was tested. In the large two branch assembly, the stent was successfully deployed in the lower arm as displayed in Fig 5E and verified by OCT imaging. Further, a stenosed module has been reopened via a separate application of a stent via balloon catheter (Fig. 5F). The full opening of the stent is visible and a widening of the stenosed area, restoring the original inner diameter of the module (Fig. 5F, SVid. 5).

3D printing is a potent way to enable generation of phantom models for the assessment of interventional medical devices^[85]^. The presented model introduces a new level of complexity compared to existing silicone tubing phantoms^[86]^. It facilitates better training of medical staff and allows for feedback of wall penetration of the higher similarity of hydrogels to native blood vessel tissue (Fig 5C, D). First studies already illustrated the benefit in training of new medical staff with 3D printed phantoms over conventional models^[28]^. Nevertheless, layer-by-layer fabrication approaches struggle to fabricate complex vascular systems and models that also encompass physiological mechanical properties of vasculature^[86][85][87]^.

Current models also lack significantly in complexity due to limitations in the manufacturing process, the material choice or the complexity of the phantom itself^[88]^. While experimental phantoms from polyurethane^[89]^ or polyvinyl alcohol^[90]^ are already showing promising results, they lack complexity compared to physiological structures. Due to its modular nature, also complex geometries can be recreated by this modular system going as far as patient specific CT data to be used to closely recreate the unique physiology via tVAM^[91]^. This could be a beneficial new system for surgeons to evaluate the situation before a surgical intervention to aid in preparation of the procedure.

The use of hydrogel-based phantom models is warranted by the materials mechanical properties that more closely recreate the physiological situation and also allow for damage to the model to be monitored, adding a new layer to medical training in interventional radiology, cardiology and vascular surgery.

Beyond the scope of this study, the switch to a biocompatible resin (for example gelatin-based materials) would allow for endothelialization and bioactive models for phantoms. Not only the transition to a natural material for the tubing would be possible, but with the option for multi-material print^[37][51]^, tVAM could help to overcome one of the current vascular phantom limitations of complexity and physiological approximation^[88]^. The addition of cells into such a biocompatible model can also enable study of the effects of endothelial damage during catheter movement or the potency of intraarterial drug release or drug-eluting stents^[88][92]^.

These are concepts that should be investigated in a dedicated study regarding the transition of the presented modular building blocks for hollow structures into functional vascular model systems.

Overall, it was demonstrated that the presented modular system can be employed for the creation of bifurcated stenosed model systems. A successful stent placement via balloon catheter has been conducted, indicating its potential as a vascular phantom for teaching or visualization purposes in medical personnel training or surgical planning.

The second model for a cell-based application set out to verify that this approach is also capable of being translated for a biological model system. It should allow for the creation of a large vascular-like network that promotes monolayer formation of endothelial cells and opens up the options of functional biological assays, for example study of fluid dynamics on cells in the lumen of the tube.

To do so, the same CAD design of modules previously presented have been fabricated with GelMA as a biocompatible resin option. A formulation with GelMA (5 % (w/w), Type A, DoF: 70-90 %) was used since it is well established for tVAM^[37][10]^. Initial seeding experiments were done in straight modules with a cell line (C166, mouse) to verify good cell attachment and growth. Both a live dead and actin staining (Fig. 6A, SVid. 6) showcased good coverage of the luminal side of the construct after 7 days of static culture in a well plate.

**Figure 6:**
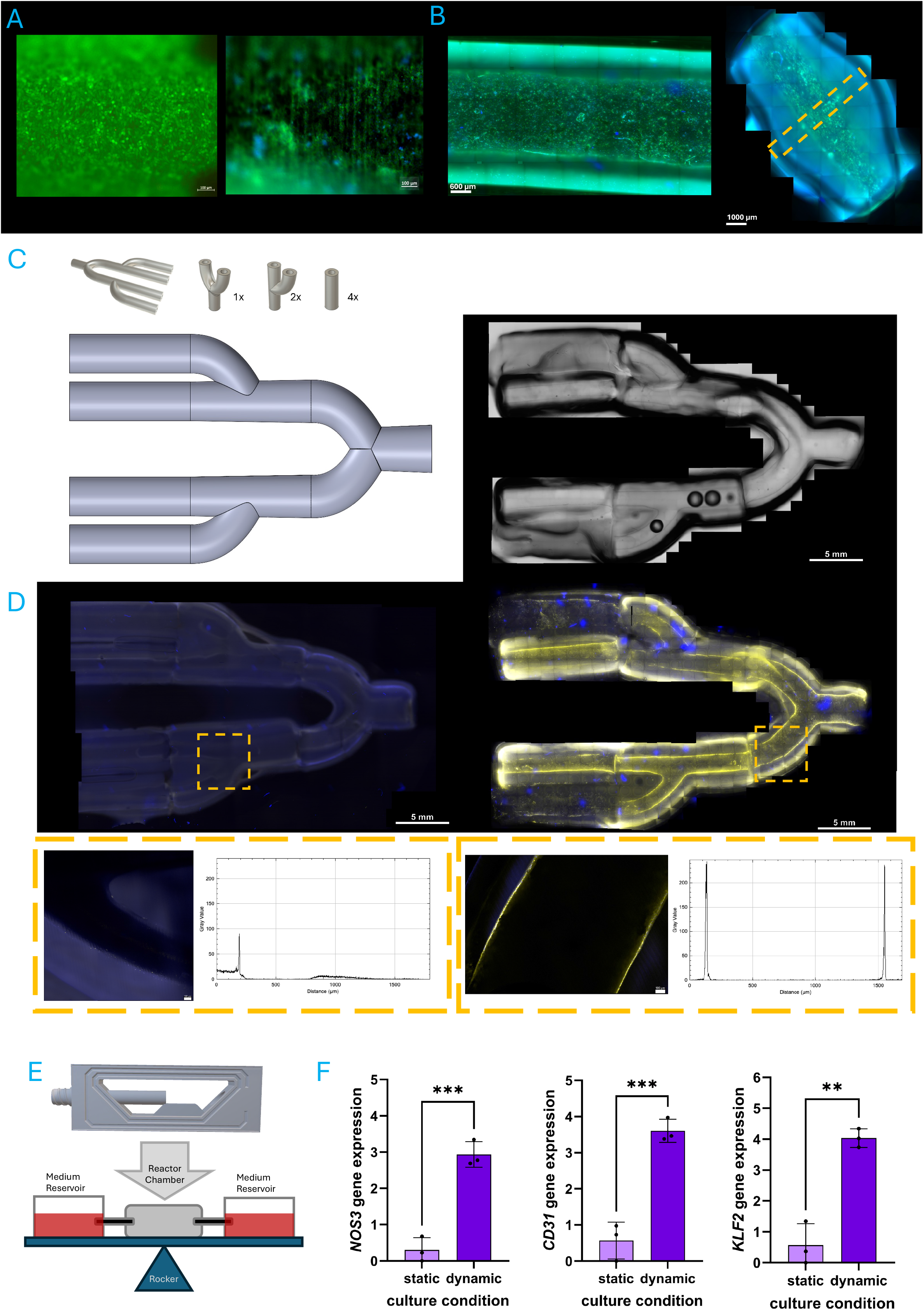
Biological evaluation of modular tube modules and assemblies out of GelMA. A) Modular module cultivated with endothelial mouse cell line after 7 days of culture, left calcein staining (green), right actin (green) and nucleus (blue) staining (SB = 100 µm). B) Modular module cultivated with HUVEC cells after 7 days of culture. Left: Actin (green) and nucleus (blue) staining (SB = C00 µm). Right, actin (green) and nucleus (blue) staining of an assembled module. Orange highlighted area indicates connection point (SB = 1 mm). C) Seven module vascular tree assembly as CAD model and printed and assembled model (SB = 5 mm). D) Assembled vascular trees with left: post-seeding of assembly and right: pre-seeding in single modules of HUVEC cells after 10 days of culture. Staining of acting (yellow) and nucleus (blue). Yellow highlight corresponds with the magnified area below where a line profile of the actin signal was performed which is shown in the intensity graph next to it (SB = 5 mm). E) Sketch of rocking perfusion setup with the custom 3D printed reactor chamber CAD scheme. F) PCR graphs of gene expression of HUVECS cultured in static and dynamic culturing conditions for NOS3, CD31 and KLF2 respectively.

For an application as a human tissue model system, human umbilical vein endothelial cells (HUVEC) were also tested and stained for actin, indicating a similarly high seeding and growth efficiency (Fig. 6B, SVid. 6). It was further evaluated if HUVECs were influenced by the connection sites between modules. A combined straight module out of two short pieces was seeded with HUVECs and evaluated if the merging area displayed any disturbances in the monolayer via actin staining (Fig 6B SVid. 7). No disturbances in the monolayer could be recorded via confocal imaging, neither in the transmission channel, nor in the fluorescent channel (Fig 6B, SVid. 6,7).

Cells from both organisms that were used in this experiment have a high relevance in current vascular model systems. Mouse cells see application in assembled models of different organs and associated pathologies^[93][94]^. HUVECS on the other hand are routinely used as a source of primary endothelial cells for many vascular model systems^[95][96][97]^. The confirmation that both types of cells work with the system confirms its applicability as a vascular model system. As results from Figure 2 showed, tVAM enables the consistent production of accurate luminal diameter modules. This capability of tVAM is highly beneficial for the generation of multi module large tubular constructs since all modules require even luminal dimensions to minimize flow turbulence and enable even cell coverage^[98][99]^. A continuous layer of cells across the connecting site also indicates that the luminal side of the constructs merge well together and allow an undisturbed monolayer formation. Further evaluation especially in the morphological differences of cells seeded on the connection site between modules will be necessary in a more functional evaluation of the system in the future.

As the presented modular system is intended to create larger tubular trees that ideally can be employed as vascular model systems, a seeding experiment of a complex vascular tree out of modular building blockss has been carried out. For this, a seven unit double bifurcated tubular tree has been designed and printed as shown in Figure 6C. The single modules were grouped in two conditions, pre- and post-seeded. For the first, each module has been seeded separately and cultured for 3 days before assembling them into the tree structure. The second condition was seeded at the same time as the first but with the tree completely assembled. The total number of cells was kept the same between all pre-seeded modules and the whole post-seeded tree. After 10 days, the samples were fixed and imaged on a fluorescent microscope to record tile scan images. The results reveal a clear endothelial lining for the pre-seeded tree that can be clearly seen in confocal detail images which have been used for a line intensity profile (Figure 6D, SFig. 5A). Overall, the monolayer formation for the pre-seeded tree was homogenous throughout the luminal area, especially on the downward facing lumen as well as the vertically oriented lumen (SFig. 5B). Compared to the post-seeded tree, the intensity of the performed actin staining was very faint and dominated by background signal due to the overall low actin signal (Figure 6D, SFig. 5C). The line profile for the post-seeded tree also implies some HUVEC presence in the wall but not as evident as in the pre-seeded tree (Figure 6D, SFig. 5B, C).

A significant challenge in vascular models is the effective endothelialization of the structure. Post seeding efficiency varies greatly by organism and geometric complexity of the structure, where large, branched ones stand out as difficult to achieve confluent monolayers^[99][100][101]^. Pre-seeding the building blocks before assembling them into a larger vascular tree facilitates homogenous cell distribution within each module that carries over into the assembly (Fig 6D, SFig. 5B). Compared to the post-assembly seeded tree, a clear difference in actin signal could be shown and confirmed by the line profile of the tube walls (Fig 6D; SFig. 5A). Similar results have been achieved by Geffert et al. in a study that employed a similar approach for the creation of modular perfusion reactors for microfluidic applications^[64]^. While their work employed fibroblasts for the cell culture experiments, the results of the presented study used vascular endothelial cells for a better transferability of the seeding results towards a vascular application.

Due to the slow growth of human endothelial cells, longer culture times would have likely improved the endothelial lining of the modules. Reports of endothelialization of simple vascular grafts mention up to 8 weeks, for complex geometries even longer^[100]^. This fact should be considered for future functional experiments carried out with this system but also supports the pre-seeding approach of single modules.

After 7 days the module was not evenly covered with ECs, predominantly on the upper part due to gravitational pull towards the bottom of the module. Even though the constructs were rotated during seeding to enhance cell attachment, the culturing after seeding was carried out in only one direction, which promoted cell accumulation in the direction of gravitational pull. A rotational culturing step would be required beyond the periodic rotation of the reactor during cell seeding to enhance an even growth of cells for future iterations^[101]^.

In this study of endothelial compatibility and growth within the modular assemblies, simple geometries and single material matrices were employed to facilitate the evaluation process. For upcoming use of this system, multi-material modules or modules with internal complications beyond simple stenosed areas will be possible. A previous study already demonstrated the flexibility with which tVAM can operate to integrate inner geometries like venous valves but also enable precise multi-material printing^[37]^, recently also shown with the assistance of computer vision^[51]^.

To ascertain the feasibility of the presented modular approach for a vascular flow model, a perfusion experiment with HUVECs in a custom DLP-printed bioreactor has been conducted. The rocking setup introduced a slow luminal liquid flow by a rocking mechanism from one reservoir into the other (Fig 6E). Single uncut modules were inserted into the reactor chamber and glued onto the connectors the same way as the modular assemblies were processed. Following assembly, they were either cultured via rocking perfusion or in a static condition to evaluate if genes related to flow (*NOS3* and *KLF2*)^[102][103]^ are expressed as well as factors related to functional endothelium (*CD31* also known as *PECAM-1*)^[104]^ (Fig. 6F). In all investigated conditions, the expression of the screeded genes was increased by up to 6 folds in perfusion conditions in comparison to static culture conditions (Fig. 6F).

Within the investigated flow system, *CD31* was significantly upregulated. This is in line with effects reported in literature for HUVECS cultured in dynamic conditions compared to static^[105] [106]^. The increase of *CD31* indicates a better cell-cell connection and barrier formation of the ECs^[104]^. Further it is also involved within the mechanosensing pathways of ECs^[107]^. Its upregulation is likely explained by the shear stress introduced by the flow within the module. In this regard, the dynamic conditions also significantly affected the expression of flow mediated factors *NOS3* and *KLF2*. *KLF2* is an integral transcription factor in multiple vascular processes including vascular tone, angiogenesis, oxidative stress and antithrombosis^[103]^. It implies that the relevant vascular pathways are activated in the tested culture conditions. Both *KLF2* and *CD31* regulate the activation of *NOS3* and by that NO synthesis, which is responsible for multiple signaling pathway interactions as well as vasodilation^[107]^.

The fact that both flow sensitive factors *KLF2* and *NOS3* are significantly upregulated is a strong indication that the perfusion of single modules is inducing expression of further flow related genes and by that likely also the corresponding protein expression in the tested HUVECs^[106]^.

Taken together, this proof of concept indicates the feasibility of modular building blocks to create a vascular perfusion model with endothelial cells that reacts to dynamic culture conditions. In future steps, larger vascular trees as well as more complex modules beyond the tested straight ones of the presented study need to be verified to maintain these favorable results. Extrapolated to the inclusion of a stenosis, since a stenosis introduces changes to flow and by that also affects mechanosensing of the seeded cells^[108][109]^. Future experiments could go deeper into this topic, potentially also with multi-material approaches in the fabrication of the module.

## Conclusion

Our results showcase that a modular approach for manufacturing large scale tubular model systems can be made via tVAM. By engineering precisely fitting modules, complex tubular systems could be manufactured that exceed the build space of commercially available tVAM systems while also reducing the required resin amount significantly. Utilizing a gluing approach to merge single modules together, a leak tight connection could be created and allowed perfusion of liquids through the lumen for both PEG and gelatine based modules. As model applications, we presented two stenosis models that were made with this modular approach. A bypass model that redirected the liquid flow around a stenosed vessel branch could recreate the flow dynamics of simulations. Further, a large model with several vessel branches enabled the placement of a balloon-catheter mounted stent in one of the stenosed modules to reopen it to its original inner diameter, effectively creating a phantom model for training purposes that better recreates tissue mechanics than routinely used systems. Biocompatibility of gelatin based modules could show a high amount of luminal cell growth of cell lines as well as HUVECS and also enabled the endothelialization of large, branched vessels by pre-seeding the single modules before connecting them. Overall, this fabrication approach allows for the recreation of many different tubular systems. This could be expanded in the future to different geometrical structures and showcases that the limitation of construct size for tVAM can be effectively mitigated, opening a new field of applications for this printing method.

## Materials and Methods

### Materials

PEGDA with number average molecular weight of 700 g mol-1, lithium phenyl-2,4,6-trimethylbenzoylphosphinate (LAP) and, if not stated otherwise, all other fine chemicals were purchased and used as is without further purification from Sigma-Aldrich. As a bioresin for VAM a selection of different materials were formulated to get a photocrosslinkable polymer precursor solution with adapted mechanical, optical, chemical and biological properties.

GelMA was synthesized according to an altered existing protocol. In short, a 10 % Gelatin solution (Type A, Sigma-Aldrich G1890, 30g) in DI-water was heated at 40 degrees until complete transparency, then at 50 °C Methacryloyl Anhydride (18 g, Sigma-Aldrich) was added. The reaction mixture was heated for 60 min, centrifuged and decanted. The product was dialyzed against di-water at 37°C (12-14kDA cutoff, Brand) for 5 days with water changes twice a day. Then the product was pooled and lyophilized in small portions.

C6-succinylated cellulose nanofibrils (C6SA-CNF) were prepared as described previously^[47]^ never-dried bleached beech sulfite dissolving pulp (Lenzing AG, Austria) was regioselectively succinylated at cellulose C6-OH with *N*-succinylimidazole, washed and fibrillated in a high-pressure homogenizer at 0.25 % wt/wt (Gaulin APV-1000, 5 passes, about 800 bar). The carboxylate content was 1.3 ± 0.1 mmol/g and determined by conductometric titration. The stock dispersion was concentrated from to 0.5 % wt/wt by evaporating water under reduced pressure (50 °C), and stored at 4 °C until use. For the CNF series, a more concentrated stock (about 1.8 wt%) was obtained by centrifugal ultrafiltration (Vivaspin 20, 100 kDa MWCO, Sartorius; 4000 × g).

### Bioresins

Photoresin containing 5 % w/w PEGDA in total was prepared by addition of the desired amount of PEGDA 700 g mol-1 to concentrated Cellulose Nanofibrils (0.5%wt, in water) and lithium phenyl-2,4,6-trimethylbenzoylphosphinate (LAP). Demineralized water was added to a final dilution of 0.24%wt CNF and 0.05 % wt/wt total LAP content. These resins were homogenized with a planetary mixer (thinky-mixer).

GelMA (5 and 8 % wt/wt) was dissolved in PBS (1x) at 37°C and 0.085% wt/wt LAP was added. For the PEGDA series, the PEGDA content was varied from 2.5 to 12.5 wt% at constant CNF (0.24 % wt/wt) and LAP contents. For the CNF series, resins with 0.08, 0.24, 0.48, 0.72 and 0.96 % wt/wt CNF were prepared at 5 wt% PEGDA from the more concentrated CNF stock. A reference resin contained 1 wt% xanthan gum instead of CNF.

### Volumetric printing

The different resin compositions were tested with a commercially available volumetric printer system (Tomolite v2, readily3d) using 10, 15 and 20 mm diameter quartz printing vials. In short, the vials were loaded with 6mL premixed bioresin each and stored at 5°C in the fridge prior to use to prevent light exposure. STL files were designed for the experiments using CAD software (Fusion360) and loaded in the Apparite software controlling the printer.

### Photorheology

A MCR 702 MultiDevice (Anton Paar, Austria) was employed for rheological measurements in conjunction with a hoenle bluepoint 4 UV curing system (Hoenle AG, Germany) which was connected for UV irradiation. The UV lamp was used in combination with a filter to ensure light exposure of the sample with wavelengths of 395 - 400 nm with an intensity of 60 %.

*Compression hydrogel tests*: Compression tests were performed with cylindric hydrogel samples with a diameter and height of 5 mm. A uniaxial tensile testing machine (BOSE, ElectroForce 5500 system, Framingham, USA) with custom sample holders and a 250 g loadcell and a 22 N loadcell was used for running and recording the experiment. The 22N cell was not sensitive enough to record the softest condition of hydrogel (PEGDA 5 wt%) which is why a switch to a more sensitive cell was necessary. A test speed of 0.015 mm/s with a total compression of 30 % (1.5 mm) was used as testing parameters. The Young’s modulus of the samples was calculated between 2.5 % and 7.5 % strain. 7 samples were measured per condition.

### OCT imaging

Optical coherence tomography (OCT Atria, Thorlabs, Germany) was used with the ThorImageOCT software (v. 5.8.4) for volumetric imaging of the printed constructs. The samples were imaged using an LK04 objective. Initially, the sample was scanned in 2D to optimize the acquisition parameters and identify and focus the region of interest. The same settings were then used for the 3D acquisition.

### Diffusion through cut and uncut hydrogels

A custom inlay system similar to trans wells was 3D printed with an DLP Printer (SL1S Speed, Prusa, Czech Republic) to accommodate hydrogel samples in the shape of discs (6 mm diameter, 1 mm thickness) to controllably measure diffusion through the hydrogel sample. To assess the leak tightness of the cut and glued approach presented in this study, uncut discs were compared to discs that were cut in half and glued together in the same fashion as the tubular modules and placed within the inlay. The inlay was then placed within a 24 well plate with the lower compartment filled with PBS containing FITC-Dextrane at 0.2 wt% (10 and 500 kDa, Merck, Darmstadt) and the upper compartment filled with only PBS. At the timepoint (0, 10, 30, 60, 120, 240 and 360 min), samples of 50 µl were taken out of the upper compartment, substituted and stored for quantification. Fluorescence was quantified by using the PBS within the well plate via a microplate-reader system (Tecan Spark, Tecan, Switzerland).

### Stent Placement and penetration assay

To help secure a large modular assembly and mimic the construction of a commonly used phantom model, the assembly has been encased in clear silicone elastomer (Sylgard 184, Dow Chemical, USA). The curing has been done according to manufacturer instructions. In a second step, a different silicone resin has been used (Dubisil 20, Dreve, Germany) to encase the clear silicone assembly and provide access via an opening created by a mold piece. The balloon catheter with a self-expanding stent out of expired medical supplies was then introduced into the modular assembly and navigated through the model before expanding the balloon using a balloon catheter pump. To test the penetration of modular modules, only the outer silicone holder has been produced for inserting a single module and connected via a custom 3D printed connector to ensure a leak tight fit. The silicone holder was filled with PBS, and the module has been closed at the endpoints with photoresin. A red dye (Allura Red AC, Sigma-Aldrich, USA) has been filled to act as a color indicator that releases when the hydrogel structure is structurally damaged. An unused balloon catheter with self-expanding stent has been introduced into the lumen of the construct and pushed into the module wall to introduce a breach, which was recorded via camera as a video (Nikon Z6, Nikon, Japan). To quantify the force to penetrate both a commonly used silicone tubing for phantom models and a hydrogel disk, a penetration experiment has been set up at a uniaxial testing machine (Z010 Zwick, ZwickRoell, Germany). A 3D printed holder to keep the samples centered has been fabricated and mounted on the lower tool port of the testing machine. For the upper tool was a custom made clamp that held a balloon catheter with self-expanding stent to be moved towards the sample at a velocity of 10 mm/min. Force was recorded via a 100 N force transducer provided by the manufacturer.

### Cell culture

Mouse yolk-sac endothelial cell line (C166, ATCC, USA) were cultured in Dulbecco’s Modified Eagle Medium with high glucose (DMEM(1x)+GlutaMAX, Gibco, Thermo Fisher, USA) supplemented with 10 % FCS and 1 % Penicilin/Streptomycin (P/S) at 37 °C and 5 % CO2 within an incubator. Human umbilical vein endothelial cells (HUVEC-2, PromoCell, Germany) were cultured in endothelial growth media-2 (EGM-2, PromoCell) supplemented with 8 % FCS and 1 % P/S at 37 °C and 5 % CO2 within an incubator. For experiments, cells of both types were detached with 0.5 % trypsin+EDTA for 5 min at 37 °C and subsequently counted and seeded at the desired concentration.

### Cell seeding of modular modules

Cell densities between 900k and 1.5 million cells per ml were used depending on experiment and cell type. Prior to seeding cells, modules were treated with a solution of fibronectin (0.2 mg/ml in PBS) for 1 h at 37 °C. After rinsing with fresh medium, each module was filled with 250 µl of cell suspension either within a well plate or mounted inside of the custom bioreactor chamber. After 40 minutes, the construct was turned around by 180 degrees to ensure cell attachment on the opposing luminal side from the initial seeding orientation. After further 40 min, the construct was turned by 90 degrees, followed by another 180 degrees turn to enable cells to attach in the four cardinal directions. Finally, the reactor chamber was filled with medium (approximately 4 ml) and sealed off with lids and one syringe filter as gas exchange valve. Well plate cultured constructs were filled with medium after seeding procedures similar to reactor cultured modules. Both well plate cultured constructs and reactor cultured modules were kept @ 37 °C and 5 % CO2.

### Gene expression analysis

To characterize the endothelial cell-like function under static and dynamic conditions, gene expression analysis was performed. After the culture period indicated, total cellular RNA isolation was performed using TRIzol (ThermoFisher Scientific, Waltham, USA) according to the manufactureŕs protocol. Briefly, the samples were lysed in 500 µl TRIzol, then chloroform was added, and the mixture was centrifuged to separate the phenol-chloroform, inter-, and aqueous phase. The aqueous phase containing the RNA was transferred into a fresh tube and precipitated with isopropanol, washed twice with 75 % ethanol, and reconstituted in RNase-free water. The RNA was precipitated with isopropanol, washed twice with 75 % ethanol, and reconstituted in RNase-free water. The RNA concentration was measured using NanoDrop™ (ThermoFisher Scientific, Waltham, USA) and up to 1000 ng RNA were transcribed into cDNA using the high-capacity cDNA reverse transcription kit (ThermoFisher Scientific, Waltham, USA) according to the manufactureŕs protocol with 0.5 µl reverse transcriptase per reaction. For the real-time polymerase chain reaction (RT-PCR) performed with the Master Mix SYBR Green® (Applied Biosystems, ThermoFisher Scientific, Waltham, USA) and ViiA7 (Applied Biosystems, ThermoFisher Scientific, Waltham, USA), 1 μl cDNA with a concentration of 2 ng/µl corresponding to the original RNA concentration were used. Primers were obtained from Biomers.net GmbH (Ulm, Germany) with exception of hRplp0 (Microsynth AG, Balgach, Switzerland) and used at 0.5 pmol/µl. The fold changes in gene expression were determined after normalization to a static control sample and the RNA amount used and pictured in bar charts with single values, mean, and standard deviation in the GraphPad Prism 10 software.

### Statistical analysis

Statistical analysis of measurements and line profiles were recorded in FIJI and processed in OriginPro 2020 (OriginLabs, USA). Significance was determined by t-test and Bonferroni-Correction. P < 0.05 is indicated by *, p < 0.01 is indicated by ** and p < 0.001 is indicated by ***.

The analysis of difference of the obtained gene expression data was performed with an unpaired t-test. p < 0.05 was considered statistically significant. Normal distribution required for performing a t-test was tested with the Shapiro-Wilk test. All statistical analyses were performed using GraphPad Prism 10 (Dotmatics, USA).

## Acknowledgements

MBK and IG contributed equally. The authors would like to thank Florian Hofmann for his support calculating computational fluid dynamics and Camilla Mussoni for proofreading and improve language. M.B.K., and T.J. thank the European Union for support on printing strategies (European Fund for Regional Development—EFRE Bayern, Bio3D-Druck project 20-3400-2-10). The DFG supported the project with a “State Major Instrumentation Programme” (INST 105022/58-1 FUGG) that enabled SEM analysis of the samples. Further, the DFG (DFG, German Research Foundation; Project No. 326998133) also provided funding for N.C.C in the framework of Collaborative Research Centre Transregio 225 (TRR 225; subproject B04; funding period 2022-2025: Prof. Süleyman Ergün and Prof. Jürgen Groll; funding period 2026-2029: Prof. Florian Kleefeldt and Prof. Jürgen Groll).

## Data Availability Statement

The data that support the findings of this study are available from the corresponding author upon reasonable request.

Figure Collection For:

