## Supplemental Figures for "Volumetric building blocks: A simple and modular approach to create perfusable models via assembly of volumetrically printed subunits"

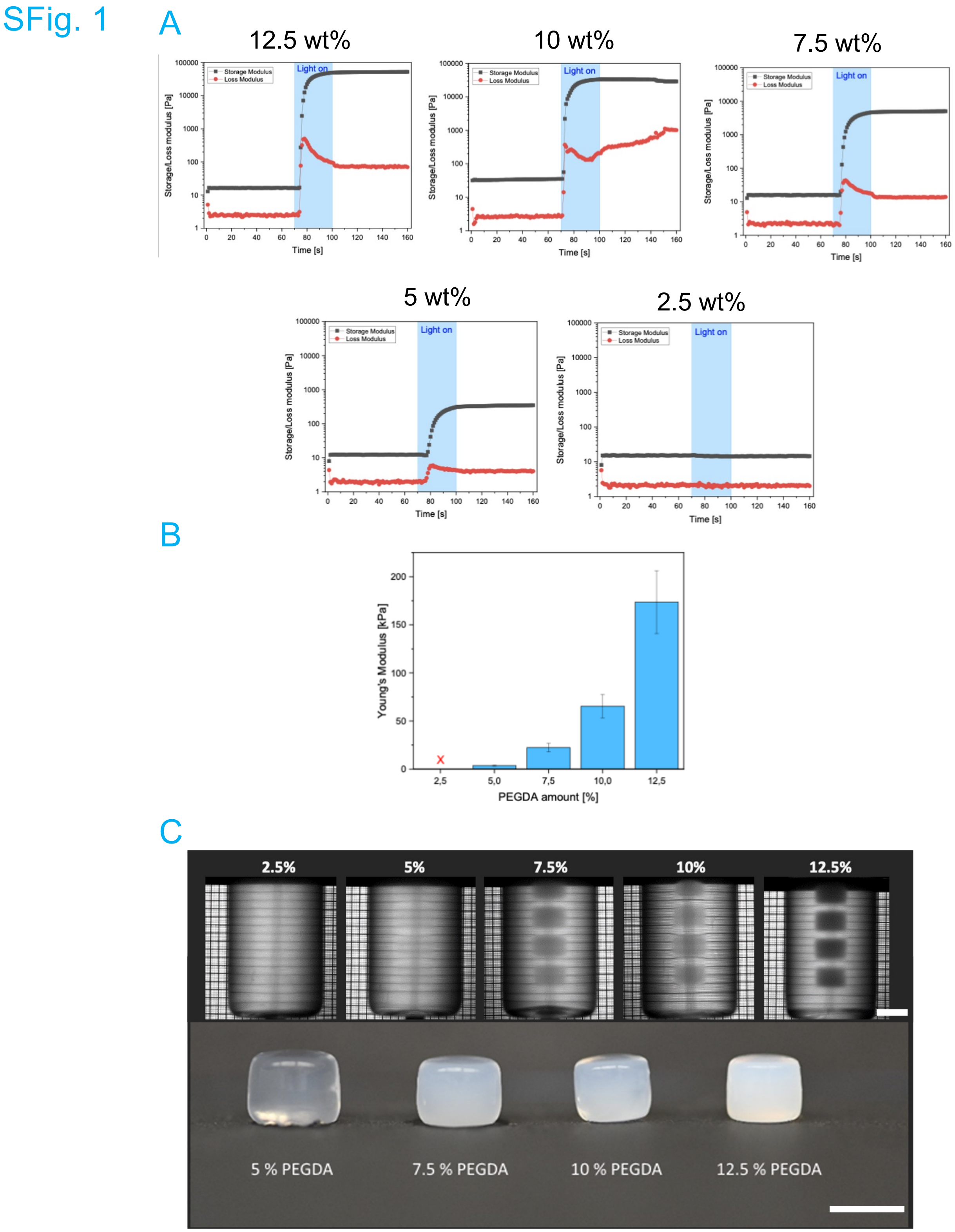


*SFig. 1: Evaluation of tVAM processability of 700 $M_n$ PEGDA. A) Photorheological behavior of PEGDA. Blue areas indicate exposure to UV light to initiate crosslinking. B) Compressive Young's modulus of different wt% of 700 $M_n$ PEGDA (n = 8). C) tVAM test prints of cylinders with increasing wt% of 700 $M_n$ PEGDA. SB = 5 mm*

#### SFig. 2

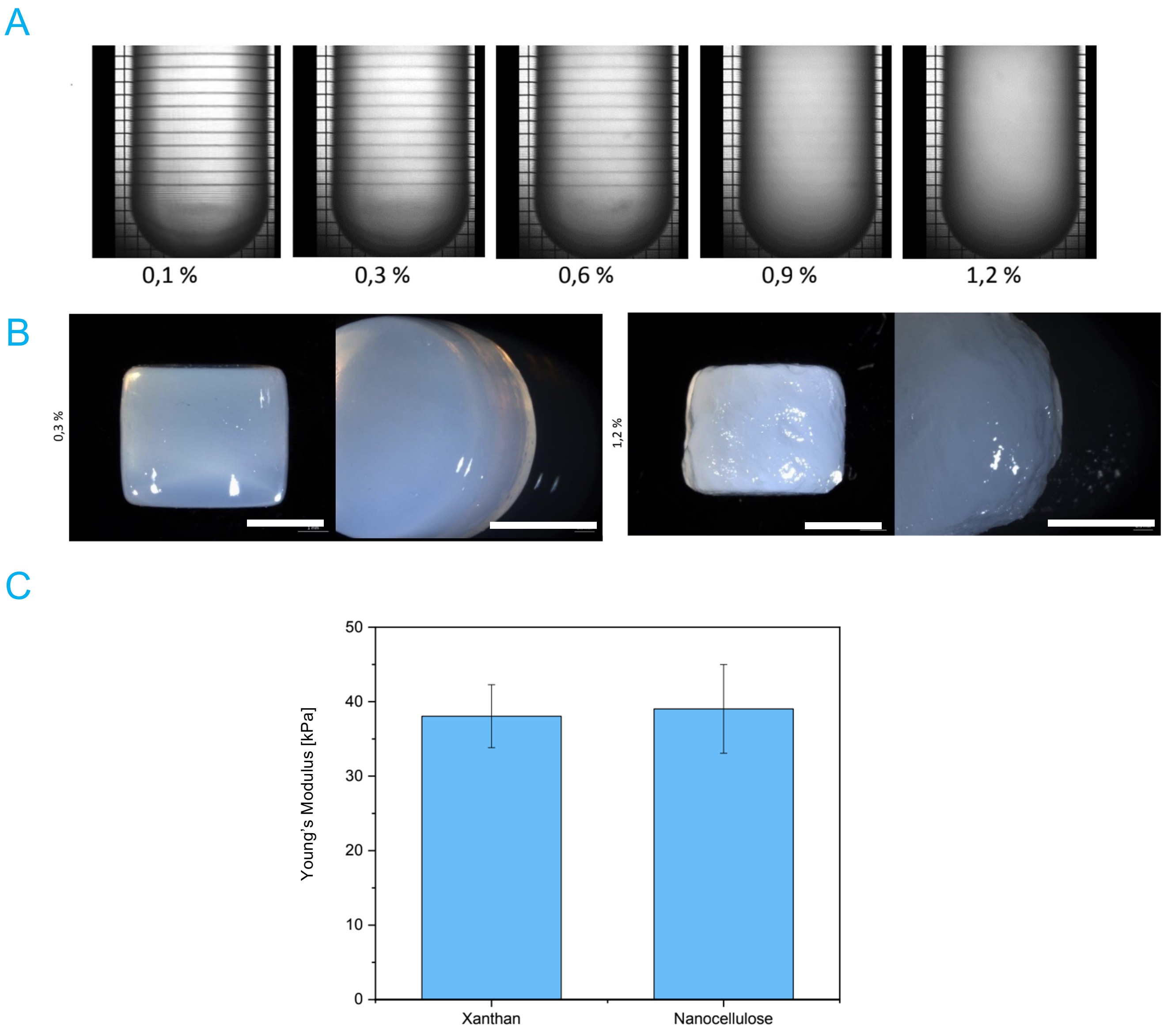


*SFig. 2: Effect of CNF on 5 wt% PEGDA hydrogels processed with tVAM. A) Increase of turbidity of PEGDA resin with increasing wt% of CNF (0.1 – 1.2 wt%) within the printing chamber of the printer. B) morphological changes of tVAM printed cylinders with CNF incorporation (0.3 wt% left and 1.2 wt% right) into 5 wt% PEGDA. SB = 2.5 mm C) Difference in Young's Modulus of 5 wt% PEGDA hydrogels modified with either Xanthan gum or CNF as thickener.*

#### SFig. 3

A

| 1 mm inner diameter | | | | | |
|---|---|---|---|---|---|
| | Design | Print | | | |
| | | M1 | M2 | M3 | Mean value |
| Inner diameter [mm] | **1.00** | 1.07 ± 0.02 | 1.03 ± 0.03 | 0.95 ± 0.02 | **1.02 ± 0.02** |
| Outer diameter [mm] | **3.00** | 3.15 ± 0.04 | 3.21 ± 0.05 | 3.25 ± 0.04 | **3.20 ± 0.04** |
| Wall thickness [mm] | **1.00** | 1.04 ± 0.04 | 1.09 ± 0.06 | 1.15 ± 0.03 | **1.09 ± 0.04** |
| **2 mm inner diameter** | | | | | |
| | Design | Print | | | |
| | | M1 | M2 | M3 | Mean value |
| Inner diameter [mm] | **2.00** | 2.07 ± 0.03 | 2.04 ± 0.05 | 2.05 ± 0.04 | **2.05 ± 0.04** |
| Outer diameter [mm] | **4.00** | 4.47 ± 0.07 | 4.48 ± 0.07 | 4.46 ± 0.07 | **4.47 ± 0.07** |
| Wall thickness [mm] | **1.00** | 1.18 ± 0.06 | 1.20 ± 0.07 | 1.18 ± 0.05 | **1.19 ± 0.06** |
| **3 mm inner diameter** | | | | | |
| | Design | Print | | | |
| | | M1 | M2 | M3 | Mean value |
| Inner diameter[mm] | **3.00** | 3.02 ± 0.04 | 3.01 ± 0.06 | 3.00 ± 0.04 | **3.01 ± 0.05** |
| Outer diameter [mm] | **5.00** | 5.69 ± 0.09 | 5.68 ± 0.11 | 5.60 ± 0.09 | **5.66 ± 0.10** |
| Wall thickness [mm] | **1.00** | 1.30 ± 0.08 | 1.31 ± 0.08 | 1.27 ± 0.07 | **1.29 ± 0.08** |

*SFig. 3: TVAM print fidelity of tubular modules compared to their CAD design.*

### SFig. 4

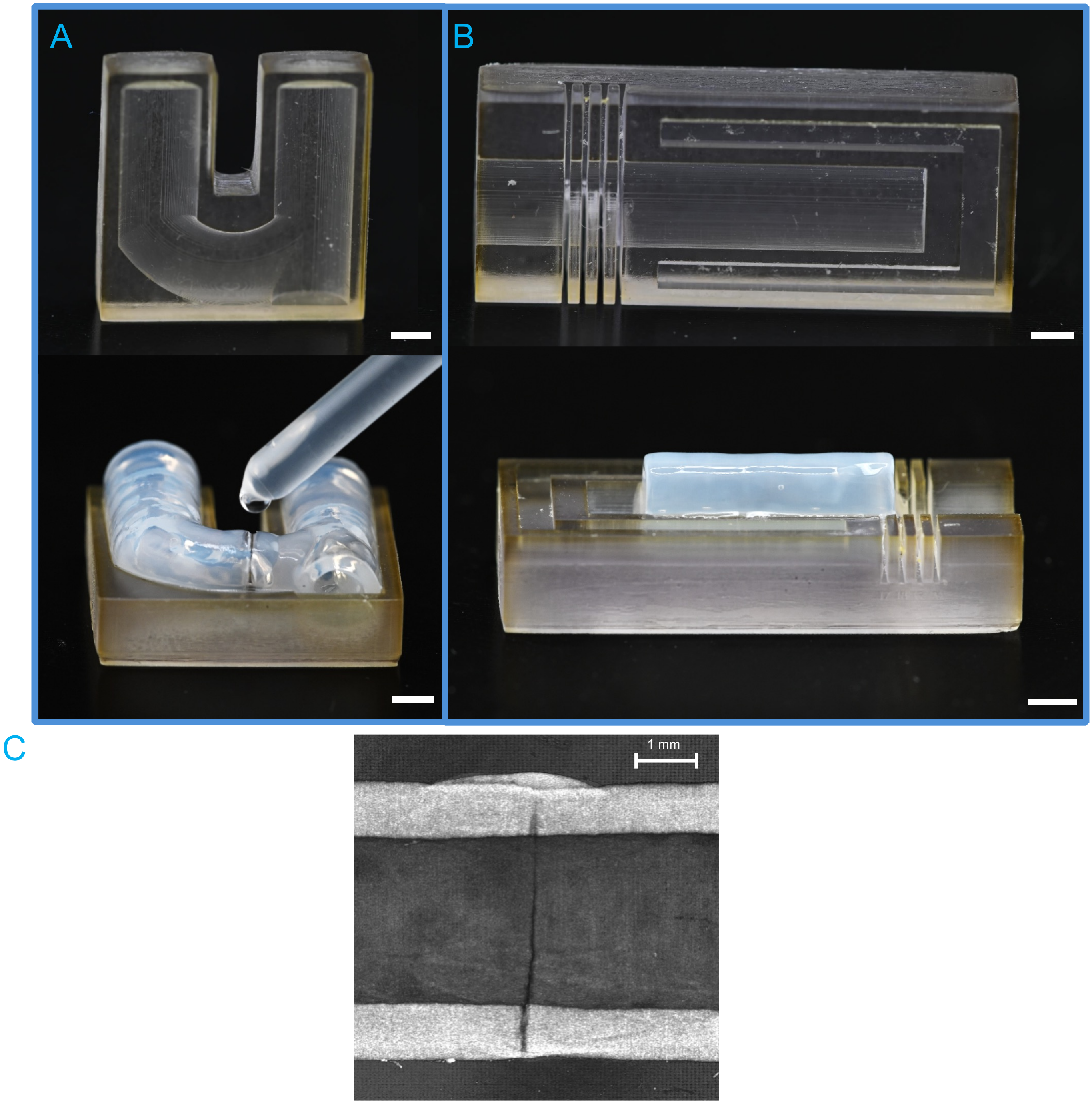


*SFig. 4: Assembly of tVAM printed modules. A) Showcase of workflow to join two modules together via joining them at the connection sites. A drop of resin at the connection sites is sufficient to create a connection. Specialized DLP printed molds help with alignment of single units. SB = 2 mm B) Specialized DLP printed mold to trim the edges of a module for easier joining of two modules. The grooves are cutting guides for a scalpel or comparable cutting tool. SB = 2 mm C) OCT image of a joined module. A small amount of resin is sufficient for a tight connection. On the upper part of the connection site, a small bulb of gluing resin is still visible. SB = 1 mm*

### SFig. 5

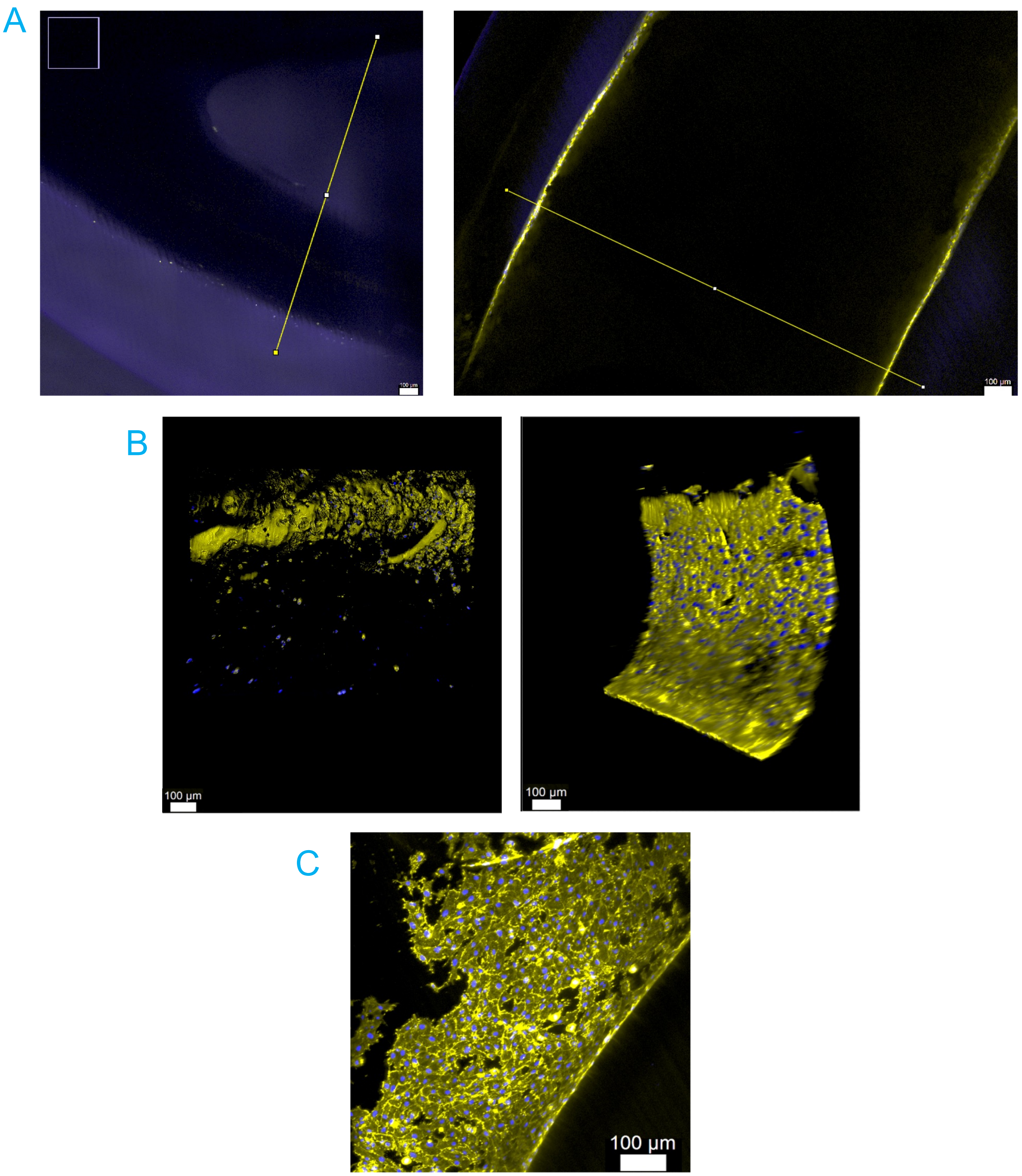


*SFig. 5: Cell growth inside of GelMA modules. A) Intensity profiles created to demonstrate cell growth on post-seeded (left) and pre-seeded (right) modules in Figure 6. SB = 100 µm B) Representative Z-stack of the image regions shown in A. Left post-seeded and right pre-seeded modules. Shown is actin (yellow) and nuclei (blue) of HUVEC cells. SB = 100 µm C) Maximum projection of a Z-Stack of a pre-seeded module. Shown is actin (yellow) and nuclei (blue) of HUVEC cells. SB = 100 µm*
