## Supplementary figures and images for "Volumetric building blocks: A simple and modular approach to create perfusable models via assembly of volumetrically printed subunits"

### Supplemental Video 6

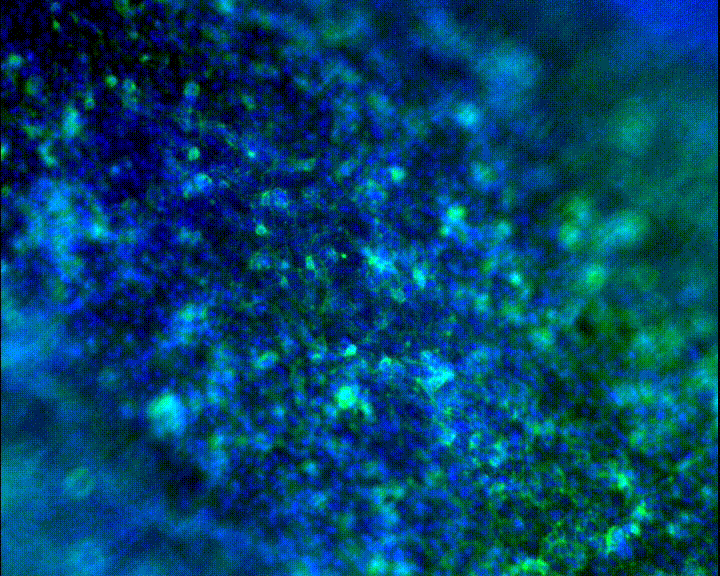

### Supplemental Video 7

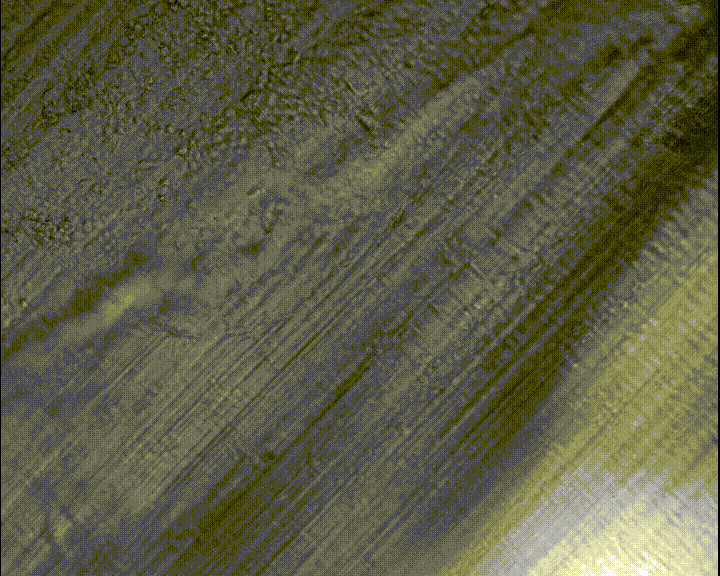
